# Functionally distinct resident macrophage subsets differentially shape responses to infection in the bladder

**DOI:** 10.1101/2020.04.18.048074

**Authors:** Livia Lacerda Mariano, Matthieu Rousseau, Hugo Varet, Rachel Legendre, Rebecca Gentek, Javier Saenz Coronilla, Marc Bajenoff, Elisa Gomez Perdiguero, Molly A Ingersoll

## Abstract

Resident macrophages are abundant in the bladder, playing key roles in immunity to uropathogens. Yet, whether they are heterogeneous, where they come from, and how they respond to infection remain largely unknown. We identified two macrophage subsets in mouse bladders, MacM in the muscle and MacL in the lamina propria, with distinct protein expression and transcriptomes. Using a urinary tract infection model, we validated our transcriptomic analyses, finding that MacM macrophages phagocytosed more bacteria and polarized to a more anti-inflammatory profile, whereas the MacL subset died rapidly during infection. During resolution, monocyte-derived cells contributed to tissue-resident macrophage pools and both subsets acquired transcriptional profiles distinct from naïve macrophages. Depletion of these altered macrophages resulted in the induction of a type 1 biased immune response to a second urinary tract infection, improving bacterial clearance. Our study uncovers the biology of resident macrophages and their response to an exceedingly common infection in a largely overlooked organ, the bladder.

## Introduction

Tissue-resident macrophages regulate immunity and are pivotal for development, homeostasis, and repair (Wynn et al., 2013). Major research efforts have uncovered roles for tissue-resident macrophages during infection, insult, and repair. However, in many cases, these studies disproportionally focus on certain organs in animals, while disregarding tissue macrophages in other locations (Epelman et al., 2014b). Importantly, macrophage function is shaped by their tissue of residence and the local environment and specific phenotypes may not be universally applicable to all tissues (Amit et al., 2016). Notably, the bladder is generally overlooked in macrophage studies, and thus, the function, origin, and renewal of bladder-resident macrophages in health and disease are poorly characterized or even completely unknown (Ingersoll and Albert, 2013; Lacerda Mariano and Ingersoll, 2018).

Tissue-resident macrophages in adult organisms originate from embryonic progenitors, adult bone marrow (BM), or a mixture of both (Epelman et al., 2014a; Ginhoux et al., 2010; Guilliams et al., 2013; Hashimoto et al., 2013; Hoeffel et al., 2015; Schulz et al., 2012; Yona et al., 2013). During development, hematopoiesis begins in the yolk sac, giving rise to erythrocytes and macrophages directly and erythro-myeloid progenitors (EMPs) (Ginhoux and Guilliams, 2016; Gomez Perdiguero et al., 2015; Hoeffel et al., 2015). As hematopoiesis declines in the yolk sac, an intra-embryonic wave of definitive hematopoiesis begins in the aorta-gonad-mesonephro, creating hematopoietic stem cells (HSCs). EMPs and HSCs colonize the fetal liver to give rise to fetal liver monocytes, macrophages, and other immune cells, whereas only HSCs migrate to the BM to establish hematopoiesis in post-natal animals (Hoeffel and Ginhoux, 2018). Embryo-derived macrophages can either self-maintain and persist into adulthood or undergo replacement by circulating monocytes at tissue-specific rates. For example, a majority of macrophages in the gut are continuously replenished by BM-derived cells, whereas brain macrophages, or microglia, are long-lived yolk sac-derived cells that are not replaced in steady state conditions (Bain et al., 2014; De Schepper et al., 2019; Ginhoux et al., 2010; Gomez Perdiguero et al., 2015). In certain conditions, origin influences macrophage behavior, for example following myocardial infarction, in which embryonic-derived cardiac macrophages promote tissue repair, whereas BM-derived macrophages induce inflammation (Dick et al., 2019). However, macrophage functions are also imprinted by their microenvironment(Gosselin et al., 2014; Lavin et al., 2014). In the small intestine macrophages in the muscle express higher levels of tissue-protective genes, such as *Retnla*, *Mrc1*, and *Cd163* compared to lamina propria macrophages, although both originate from adult BM (Gabanyi et al., 2016).

While the origin and maintenance of bladder-resident macrophages is currently unknown, these macrophages do play a role in the response to urinary tract infection (UTI), which impacts up to 50% of all women at some point in their lifetimes (Foxman, 2002; Lacerda Mariano and Ingersoll, 2018). The immune response to *uropathogenic Escherichia coli* (UPEC) infection in the bladder is characterized by robust cytokine expression leading to rapid infiltration of large numbers of neutrophils and classical Ly6C^+^ monocytes (Hang et al., 1999; Haraoka et al., 1999; Ingersoll et al., 2008; Mora-Bau et al., 2015; Shahin et al., 1987; Zychlinsky Scharff et al., 2019). Although essential to bacterial clearance, neutrophil and monocyte infiltration likely also induce collateral tissue damage. Targeted depletion of these cell types individually is associated with reduced bacterial burden after primary infection in mice, whereas elimination of both cell types together leads to unchecked bacteria growth (Haraoka et al., 1999; Ingersoll et al., 2008; Mora-Bau et al., 2015). Tissue-resident macrophages also take up a large number of bacteria during UTI, however, transient depletion of resident macrophages just before infection does not change bacterial clearance in a first, or primary UTI (Mora-Bau et al., 2015). Remarkably, the absence of macrophages in the early stages of a primary UTI significantly improves bacterial clearance during a second, or challenge, infection (Mora-Bau et al., 2015). Exactly how the elimination of resident macrophages improves the response to a challenge infection is unclear, particularly as tissue-associated macrophages return to homeostatic numbers in the time interval between the two infections. Of note, improved bacterial clearance is lost in macrophage-depleted mice that are also depleted of CD4^+^ and CD8^+^ T cells, suggesting that macrophages modulate T cell activation or limit differentiation of memory T cells, as observed in other tissues (Bergsbaken et al., 2017; Denning et al., 2007; Goplen et al., 2019; Panea et al., 2015; Taylor et al., 2006). For example, ablation of embryonic-derived alveolar macrophages results in increased numbers of CD8^+^ resident memory T cells following influenza infection in mice (Goplen et al., 2019). In the gut, monocyte-derived macrophages support the differentiation of CD8^+^ tissue-resident memory T cells by production of IFN-β and IL-12 during *Yersinia* infection (Bergsbaken et al., 2017). Indeed, the opposing roles of macrophages in modulating T cell responses in the lung and gut support that tissue type and/or ontogeny determine how macrophages may influence adaptive immunity (Ginhoux and Guilliams, 2016).

To understand the role of bladder-resident macrophages, we investigated the origin, localization, and function of these cells during infection. We identified two subpopulations of resident macrophages in naïve mouse bladders with distinctive cell surface proteins, spatial distribution, and gene expression profiles. We found that bladder macrophage subsets were long-lived cells, slowly replaced by BM-derived monocytes over the lifetime of the mouse. During UTI, the macrophage subsets differed in their capacity to take up bacteria and survive infection, however, both subsets were replaced by BM-derived cells following resolution of infection. Thus, after a first infection, macrophage subsets had divergent transcriptional profiles compared to their naïve counterparts, shaping the response to subsequent UTI.

## Results

### Two spatially distinct macrophage populations reside in the naïve bladder

We reported that macrophage depletion before a first UTI improves bacterial clearance during challenge infection (Mora-Bau et al., 2015). Thus, we initiated a follow-up study to investigate the role of bladder resident macrophages during UTI. Using the macrophage-associated cell surface proteins CD64 and F4/80 (Austyn and Gordon, 1981; Gautier et al., 2012), we identified a uniformly CD64 and F4/80 double positive resident macrophage population in naïve bladders from 7-8 week old female CX_3_CR1^GFP/+^ mice. This transgenic mouse is widely used to distinguish macrophage populations in other tissues as the chemokine receptor CX_3_CR1 is expressed by monocytes and macrophages at some point in their development (Jung et al., 2000). In most tissue resident macrophages are either GFP+ as they express CX_3_CR1 or GFP negative because they no longer express CX_3_CR1 (Yona et al., 2013), therefore, we were surprised to observe heterogeneity in GFP expression levels, revealing potentially two subpopulations (Mac^lo^ and Mac^hi^) of CD64^+^ F4/80^+^ macrophages in the bladder (**Figure 1A**). The Mac^hi^ expressing population was present in significantly greater numbers and proportions compared to the Mac^lo^ population (**Figure 1A**). As CX_3_CR1 deficiency results in decreased macrophage numbers and frequency in the intestine and brain tissue, and the transgenic CX_3_CR1^GFP/+^ mouse we used is hemizygous for this receptor (Jung et al., 2000; Medina-Contreras et al., 2011; Tang et al., 2014), we investigated whether our putative bladder resident macrophage subsets were similarly present in wildtype C57BL/6 mice. Using the same gating strategy and an anti-CX_3_CR1 antibody, we clearly identified two distinct macrophage populations with different CX_3_CR1 expression levels in 7-8 week old naïve female wildtype mice (**Figure 1B**). Notably, wildtype mice had similar numbers and proportions of each macrophage subset (**Figure 1B**).

**Figure 1.**
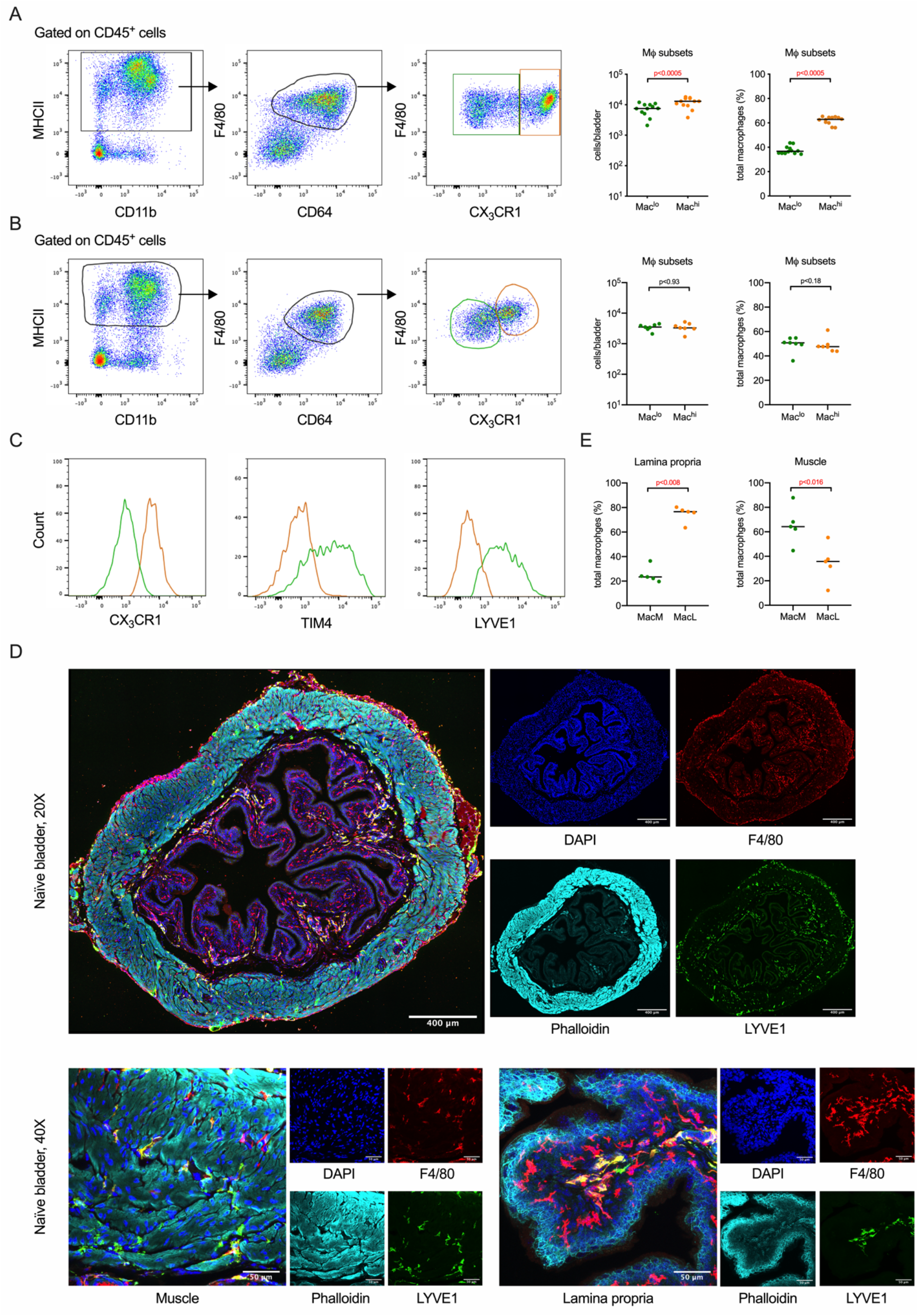
Two macrophage subsets are resident in adult naïve mouse bladders. (**A-C**) Bladders from 7-8 week old naïve female CX_3_CR1^GFP^ and C57BL6/J mice were analyzed by flow cytometry. (**A-B**) Dot plots depict the gating strategy to identify macrophages by surface protein expression, and subsets by CX_3_CR1 expression, in (**A**) CX_3_CR1^GFP^ and (**B**) C57BL6/J mice. The graphs show the total cell number on a log-scale (left) and proportion (right) of macrophages in the bladder by subset derived from cytometric analysis in (**A**) CX_3_CR1^GFP^ and (**B**) C57BL6/J mice. (**C**) Histograms show the relative expression of CX_3_CR1, TIM4, and LYVE1 on macrophage subsets in C57BL6/J mice, in which Mac^lo^ is shown in green and Mac^hi^ is in orange. (**D**) Micrographs are representative confocal images of naïve bladder from C57BL6/J mice at 20X and 40X. Merged images and single channels with the target of interest are shown. (**E**) Graphs show the proportion of each macrophage subset in the lamina propria and muscle of naïve C57BL6/J mice. Data are pooled from 3 experiments, n=3-6 mice per experiment. Each dot represents one mouse, lines are medians. Significance was determined using the nonparametric Mann-Whitney nonparametric test to compare the numbers of macrophages subsets (**A, B**) and the nonparametric Wilcoxon matched-pairs signed rank test to compare the percentages of each macrophage subset (**A, B, E**). All calculated p-values are shown, p-values <0.05, indicating statistical significance, are in red.

Next, we assessed the surface expression level of proteins known to define macrophage subsets in other tissues (Chakarov et al., 2019). We observed that the efferocytic receptor TIM4 and hyaluronan receptor LYVE1 were expressed by the Mac^lo^ population, whereas the Mac^hi^ population was TIM4 and LYVE1 negative (**Figure 1C)**. To determine the spatial orientation of the subsets, we stained naïve female C57BL/6 bladders with antibodies to F4/80 and LYVE1, and phalloidin to demarcate the muscle layer from the lamina propria (**Figure 1D**). We quantified the number of each subset in these two anatomical locations, observing a higher percentage of the Mac^lo^ macrophage subset in the muscle compared to the Mac^hi^ macrophage subset (**Figure 1E**). Macrophages in the lamina propria were predominantly of the Mac^hi^ phenotype (**Figure 1E**). Thus, the phenotypical differences we observed in bladder-resident macrophage subsets extended to differential tissue localization. Given their spatial organization, we renamed the Mac^lo^ subset “MacM” for muscle and the Mac^hi^ subset “MacL” for lamina propria. Altogether, these results reveal that two phenotypically distinct macrophage subsets reside in different regions of the naïve bladder.

### Bladder-resident macrophages, seeded from the yolk sac, are replaced by HSC-derived macrophages after birth

We next investigated whether macrophage heterogeneity in adult mouse bladders arose due to distinct developmental origins of the subsets. We analyzed bladders from newborn C57BL/6 pups by confocal imaging and CX_3_CR1-GFP expressing E16.5 embryos and newborn mice by flow cytometry. We observed that in E16.5 and newborn animals, a single CX_3_CR1^hi^ macrophage population was present in the muscle and lamina propria of the bladder, and these cells stained positively for LYVE1 in confocal images of newborn mouse bladder, as observed for many fetal tissues, supporting that diversification of bladder macrophage subsets occurs after birth (**Figure 2A**). We hypothesized that in adult mice, macrophage subsets might arise following differentiation of cells seeded from embryonic progenitors or that one subset is derived from embryonic macrophages, whereas the second subset arises from BM-derived monocytes (Mossadegh-Keller et al., 2017). To test these hypotheses, we used the *Cdh5*-CreERT2 Rosa26^tdTomato^ transgenic mouse, in which the contribution of distinct hematopoietic progenitor waves to immune cell populations can be followed temporally, such that treatment of pregnant mice with 4-hydroxytamoxifen (4OHT) at E7.5 labels yolk sac progenitors and their progeny and treatment at E10.5 labels HSC and their cellular output (Gentek et al., 2018). After treatment with 4OHT at E7.5, in which microglia were labeled as expected (Ginhoux et al., 2010; Gomez Perdiguero et al., 2015), we found a significantly higher proportion of labeled bladder macrophages in E16.5 embryos and newborn mice compared to monocytes (**Figure 2B**), which was not the case in adult (8-11 week old) mice (**Figure 2C**), indicating that yolk sac macrophages are gradually diluted and in the adult the subsets are comprised of HSC-derived macrophages, similar to that seen in the kidney (Sheng et al., 2015). In accordance with this, low levels of E10.5-labeled macrophages were detected in embryonic bladders (**Figure 2D**) and their frequency increased over time, although to a lesser degree than monocytes (**Figure 2D, E**). Of note, both subsets found in the adult bladder showed similar frequencies of E10.5 labeling (**Figure 2E**). Together, these results demonstrate that adult bladder macrophages are HSC-derived and the macrophage subsets cannot be distinguished from each other by their ontogeny.

**Figure 2.**
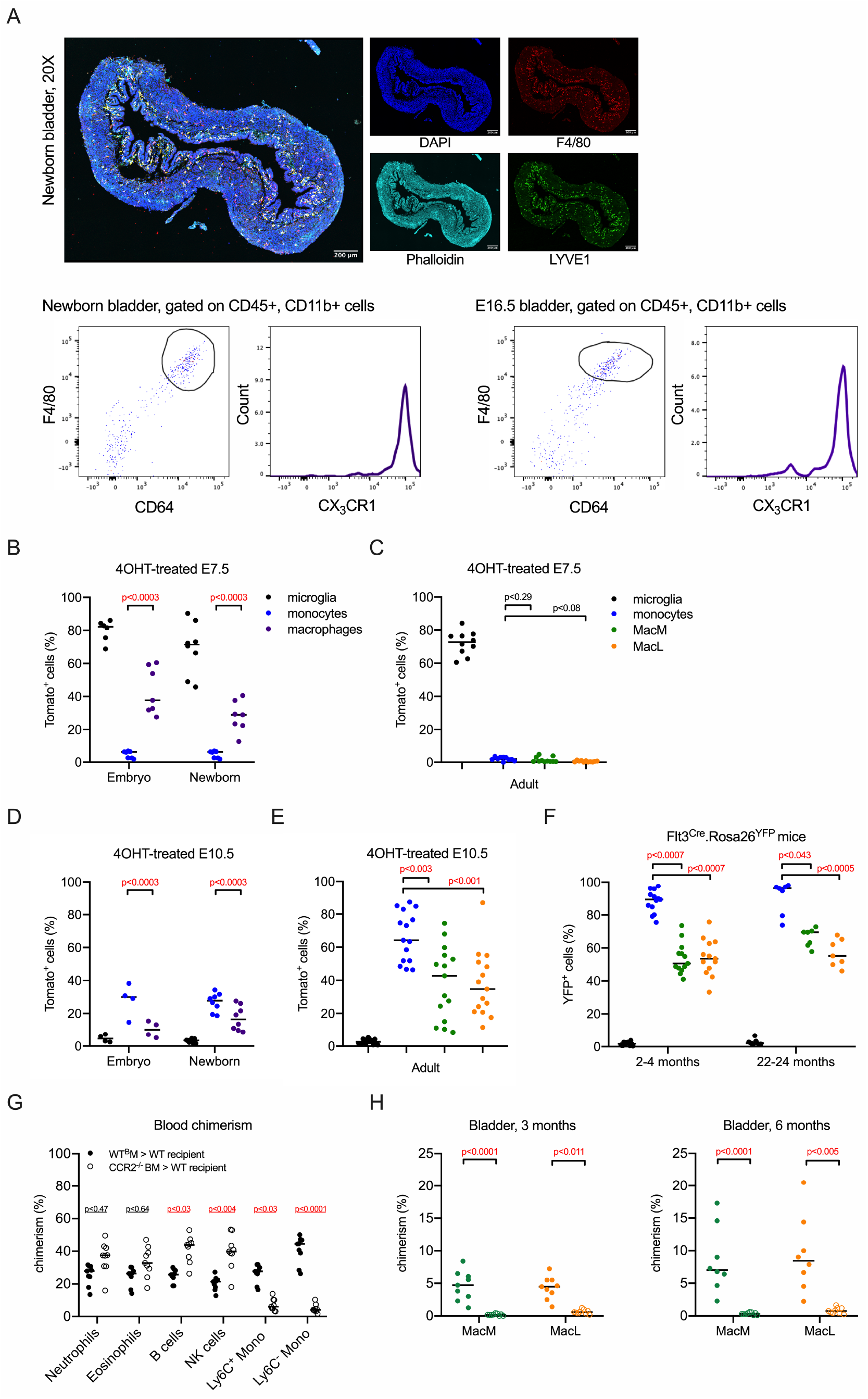
Adult bladder resident macrophages are long-lived HSC-derived cells. (**A**) Micrographs show a merged confocal image and single channels of a representative bladder from a C57BL/6 newborn mouse. Dot plots show gating strategy and CX_3_CR1 expression for macrophages in *Cdh5*-CreERT2 Rosa26^tdTomato^ CX_3_CR1^GFP^ newborn mice (left) and E16.5 embryos (right). (**B-E**) Graphs depict reporter recombination (percentage of tomato+ cells) in microglia (black dots), circulating monocytes (blue dots), total bladder macrophages (purple dots), or MacM (green dots) and MacL macrophages (orange dots) in *Cdh5*-CreERT2 Rosa26^tdTomato^ (**B**) E16.5 embryos and newborns treated with 4OHT at E7.5, (**C**) adult mice treated with 4OHT at E7.5, (**D**) E16.5 embryos and newborns treated with 4OHT at E10.5, (**E**) adult mice treated with 4OHT at E10.5. (**F**) Graph shows percentage of YFP+ cells in microglia (black dots), circulating monocytes (blue dots), MacM (green dots) and MacL macrophages (orange dots) in naïve adult male Flt3^Cre^ Rosa26^YFP^ mice. (**G-H**) Adult C57BL/6 CD45.2 mice, irradiated with a lead shield protecting the bladder, were reconstituted with CCR2^+/+^ CD45.1 BM cells and adult CD45.1 shield-irradiated mice were reconstituted with CCR2^−/−^ CD45.2 BM cells. (**G**) Graph shows percentage of donor cells in circulating leukocytes in mice transplanted with CCR2^+/+^ (filled circles) or CCR2^−/−^ BM (open circles) at 3 months after transplantation. (**H**) Graphs show percentage of donor cells contributing to bladder macrophage subsets in mice transplanted with CCR2^+/+^ (filled circles) or CCR2^−/−^ BM (open circles) at 3 and 6 months after BM transplantation. Data are pooled from 2-3 experiments, n=3-6 mice per experiment in adult mice, and n=2-4 mice per experiment with embryos and newborn mice. Each dot/circle represents one mouse, lines are medians. Significance was determined using the nonparametric Mann-Whitney tests to compare macrophages or subsets to monocytes (**B**-**F**) or CCR2^+/+^ to CCR2^−/−^ recipients (**G**-**H**), and the resulting p-values were corrected for multiple testing using the false discovery rate (FDR) method. All calculated/corrected p-values are shown and p-values meeting the criteria for statistical significance (p <0.05) are depicted in red.

To confirm that HSC-derived progenitors contribute to the bladder resident macrophage pool, we analyzed bladders from adult Flt3^Cre^ Rosa26^YFP^ mice. In this transgenic mouse, expression of the tyrosine kinase receptor Flt3 in multipotent progenitors leads to expression of YFP in the progeny of these cells, such as monocytes, whereas microglia, arising from yolk sac progenitors are essentially YFP negative (Boyer et al., 2011). Recombination rates driven by Flt3 are very low during embryonic development, but blood monocyte labelling reaches 80-90% in adult mice (Schulz et al., 2012). Therefore, if tissue-resident macrophages arise from BM-derived monocytes, labelling in adult mice should be similar to blood monocytes, whereas the presence of Flt3 negative tissue macrophages would indicate they originated from either embryonic or adult Flt3-independent progenitors. We observed that in 2-4 month and 22-24 month old mice, about 50% of each macrophage subset was YFP^+^, which was significantly lower compared to circulating monocytes (**Figure 2F**). This observation and those from the *Cdh5*-CreERT2 mice support that adult bladder macrophage subsets are derived from embryonic progenitors and HSCs, with no contribution from yolk sac macrophages. Additionally, the lack of equilibration of YFP labelling in the bladder with blood monocytes at 22-24 months suggests that tissue resident macrophages are not rapidly replaced by BM-derived macrophages in the steady state.

To determine the replacement rate of bladder-resident macrophages by BM-derived cells, we evaluated bladder-shielded irradiated mice transplanted with congenic BM from wildtype or CCR2^−/−^ mice. Monocytes depend on CCR2 receptor signaling to exit the BM into circulation (Serbina and Pamer, 2006). At 12 weeks, when engraftment was established, we observed 27.6% of circulating Ly6C^+^ monocytes were of donor origin in mice reconstituted with wildtype BM, whereas only 6% of Ly6C^+^ monocytes were of donor origin in wildtype mice receiving CCR2^−/−^ BM (**Figure 2G**). B and NK cells were replenished to a greater extent in mice reconstituted with CCR2^−/−^ BM compared to mice reconstituted with CCR2^+/+^ BM, which could be due to different engraftment efficiencies between CD45.1 and CD45.2 BM (Mercier et al., 2016; Waterstrat et al., 2010) (**Figure 2G**). In mice reconstituted with wildtype BM, 3.8% of MacM and 4% MacL were of donor origin at 3 months post-engraftment. At 6 months after irradiation, 7% of MacM and 8% of MacL macrophages were of donor origin (**Figure 2H**). By dividing the median macrophage subset chimerism (7% or 8%) by the median circulating Ly6C^+^ monocyte chimerism at 6 months (27.7%), we determined that 28% of MacM and MacL were replaced by BM-derived monocytes within 6 months. Chimerism in bladder macrophage subsets was markedly reduced in CCR2^−/−^ BM recipients, suggesting that monocytes slowly replace bladder macrophage subsets in a CCR2-dependent manner (**Figure 2H**).

Taken together, these results reveal that establishment of distinct bladder resident macrophage subsets occurs postnatally. Yolk sac macrophages initially seed the fetal bladder, but are later replaced by HSC-derived macrophages. In adult mice, bladder macrophage subsets are maintained, at least in part, though a slow replacement by BM-derived monocytes. Importantly, MacM and MacL macrophages do not differ in their developmental origin or rate of replacement by monocytes suggesting that a particular niche in adult tissue may be responsible for macrophage specialization into phenotypical and functionally distinct macrophage subsets.

### Resident bladder macrophage subsets have divergent transcriptomes

Although bladder-resident macrophage subsets had similar ontogeny, their distinct spatial localization and surface protein expression suggested that they have different functions. To test this hypothesis, we first analyzed gene expression profiles of naïve adult female MacM and MacL macrophages using bulk RNA sequencing. To formally demonstrate that our cells of interest are macrophages, we aligned the transcriptomes of the bladder macrophage subsets with the macrophage core signature list published by the Immunological Genome Consortium and the bladder macrophage core list from the mouse cell atlas single-cell database (Gautier et al., 2012; Han et al., 2018). The MacM and MacL subsets expressed 80% of the genes from the Immunological Genome Consortium macrophage core signature list and more than 95% of the genes in the bladder macrophage core list (**Figure 3A**), supporting that our cells of interest are indeed fully differentiated tissue resident macrophages.

**Figure 3.**
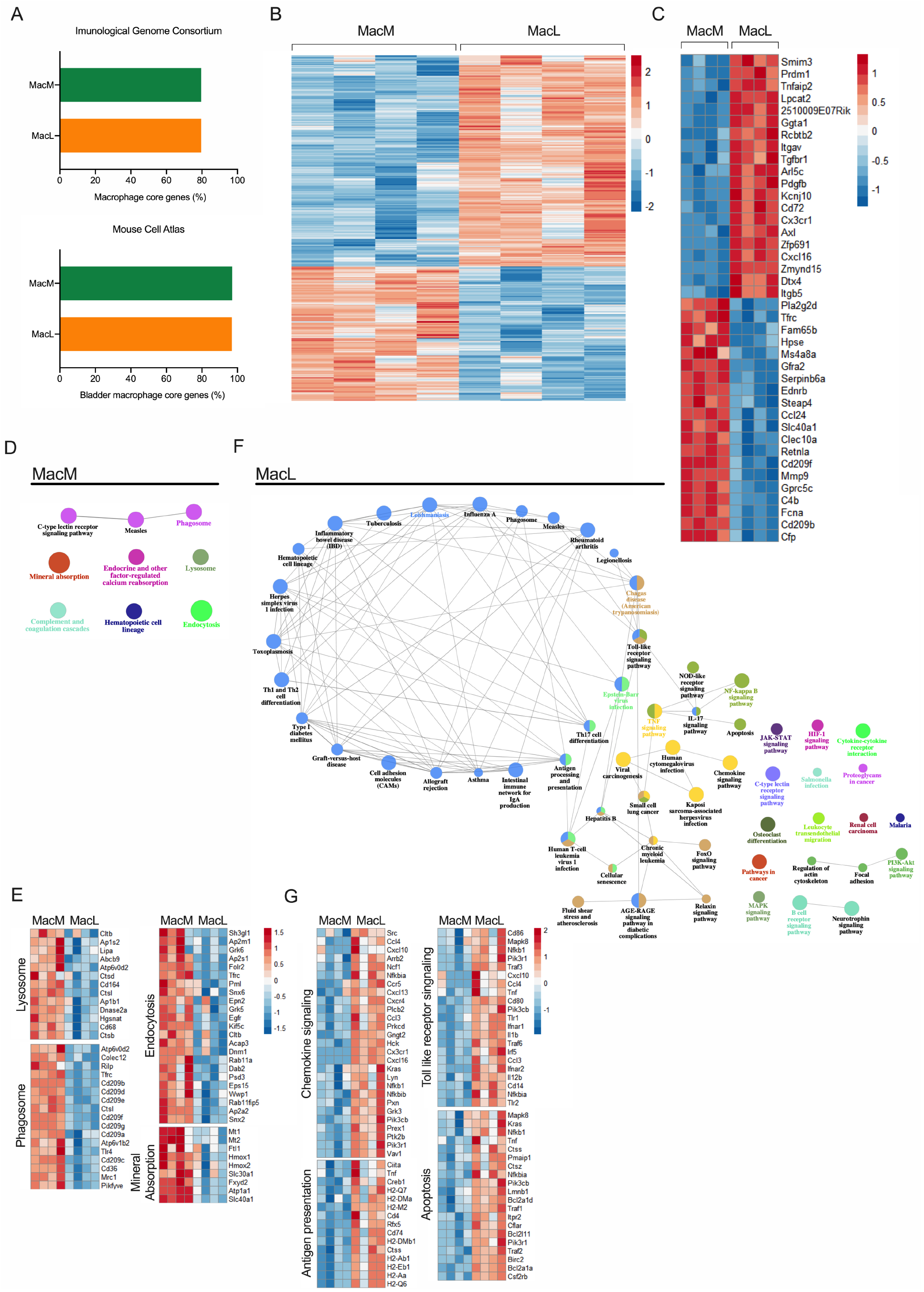
Macrophage subsets have different transcriptional programs in the naïve bladder. MacM and MacL macrophages were sorted from 7-8 week old female naïve adult C57BL/6 mouse bladders and analyzed by RNAseq. **(A)** Bar graphs show the percentage of overlap in gene expression between MacM and MacL macrophages and the macrophage core signature defined by the Immunological Genome Consortium (top) or the Mouse Cell Atlas (bottom). (**B-C**) The heat maps show the gene expression profile of the 1,475 differentially expressed genes and (**C**) the 20 most differentially expressed genes between the MacM and MacL subsets. (**D-G**) Using KEGG pathway analysis of significantly up-regulated genes, depicted are (**D**) pathways enriched in MacM macrophages, (**E**) up-regulated genes associated with selected pathways in MacM macrophages, (**F**) pathways enriched in MacL macrophages, up-regulated genes associated with selected pathways in MacL macrophages. In **D, F**, the size of the nodes reflects the statistical significance of the term. (*Q*<0.05; terms >3 genes; % genes/term >3; *κ* 0.4).

We observed that 1,475 genes were differentially expressed between naïve MacM and MacL macrophages, in which 899 genes were positively regulated and 576 genes were negatively regulated in the MacL subset relative to MacM macrophages (**Figure 3B**). In the top 20 differentially expressed genes (DEG), MacM macrophages expressed higher levels of *Clec10a*, *Retnla*, and *Ccl24*, which are associated with an alternatively-activated macrophage phenotype (Nguyen et al., 2011; Ploeger et al., 2013; Raes et al., 2005; Voehringer et al., 2007); genes involved in iron metabolism, such as *Tfrc*, *Steap4*, and *Slc40a1* (Nairz et al., 2017); and genes from the complement cascade, including *C4b* and *Cfp* (**Figure 3C**). In the same 20 most DEG, MacL macrophages expressed greater levels of *Cx3cr1*, *Cd72*, *Itgb5*, *Axl*, and *Itgav*, which are associated with phagocytosis, antigen presentation, and immune response activation (Agostini et al., 1999; Bain and Schridde, 2018; Schridde et al., 2017) (**Figure 3C**). MacL macrophages also expressed inflammatory genes, such as *Cxcl16*, a chemoattractant for T and NKT cells (Jiang et al., 2005; Matsumura et al., 2008), and *Lpcat2* and *Pdgfb*, which are involved in the metabolism of inflammatory lipid mediators (Jackson et al., 1998; Shimizu, 2009) (**Figure 3C**). Using gene set enrichment analysis of the DEG to detect pathways up-regulated in the macrophage subsets, we observed that the MacM subset expressed genes linked to pathways such as *endocytosis*, *mineral absorption*, *lysosome*, and *phagosome* (**Figure 3D**). Within the phagosome and endocytosis pathways, genes critical for bacterial sensing and alternative activation such as *Tlr4*, *Mrc1* (encoding for CD206), *Cd209*, and *Egfr* (Beutler, 2000; Montoya et al., 2009; Tang et al., 2020) were increased in the MacM subset. In the mineral absorption pathway, genes controlling iron metabolism that also enhance bacterial killing such as *Hmox1* and *Hmox2* were upregulated in MacM macrophages (Sukhbaatar and Weichhart, 2018) (**Figure 3E**). In the MacL subset, genes linked to diverse inflammatory pathways, including *Toll-like receptor signaling, apoptosis, antigen processing and presentation,* and *chemokine signaling* were present, as were many infectious and inflammatory disease-related pathways (**Figure 3F**). Within these pathways, the MacL subset expressed genes related to bacterial sensing such as *Tlr1, Tlr2,* and *Cd14*, which are important for bacterial sensing. In addition, MacL macrophages likely are involved in the initiation of inflammation, as this subset had higher expression of inflammatory cytokine genes such as *Il1b, Tnf, Ccl3, Ccl4, Cxcl10, Cxcl16,* and *Nfkb1* (**Figure 3G**).

These findings suggest that MacM macrophages are more anti-inflammatory with increased endocytic activity, which is a common feature of highly phagocytic resident macrophages (A-Gonzalez et al., 2017), and as such may play a prominent role in bacterial uptake or killing during infection. MacL, on the other hand, may play a greater role in antigen presentation and initiation or maintenance of inflammation.

### Macrophage subsets respond differently to infection

As we observed enrichment of genes belonging to *endocytosis*, *lysosome*, and *phagosome* pathways in the MacM subset, we reasoned that the macrophage subsets may differentially take up bacteria during infection. To test our hypothesis, we used a well-described mouse model of urinary tract infection, in which we transurethrally infect adult female mice via catheterization with 10^7^ counting forming units (CFU) of UPEC strain UTI89-RFP, which expresses a red fluorescent protein (Mora-Bau et al., 2015). At 24 hours post-infection (PI), we investigated bacterial uptake by macrophage subsets (**Figure 4A, B**). Despite that MacM macrophages are farther from the infected urothelium than MacL macrophages, we observed that 20% of MacM and only 10% of MacL subsets contained bacteria at 24h PI, providing functional evidence to support the transcriptional data that MacM macrophages have a superior phagocytic capacity compared to MacL macrophages (**Figure 4B**). Taking the total population of UPEC-containing macrophages, we observed that ~80% of these cells belonged to the MacM subset, whereas the MacL subset comprised only 20% of this population, which was surprising given that MacM and MacL exist in the bladder in a 1:1 ratio (**Figure 4B, Figure S1A**). We also measured polarization of the macrophage subsets in naïve and infected bladders by analyzing the expression of IL-4Rα by flow cytometry (**Figure 4C**). IL-4Rα is the receptor of IL-4 and IL-13, two cytokines that drive alternative activation in macrophages (Gordon and Martinez, 2010). Both macrophage subsets had increased expression of IL-4Rα at 24h PI compared to their naïve counterparts; however, MacM macrophages had consistently higher expression levels of IL-4Rα compared to MacL macrophages in naïve or infected tissue (**Figure 4C**).

**Figure 4.**
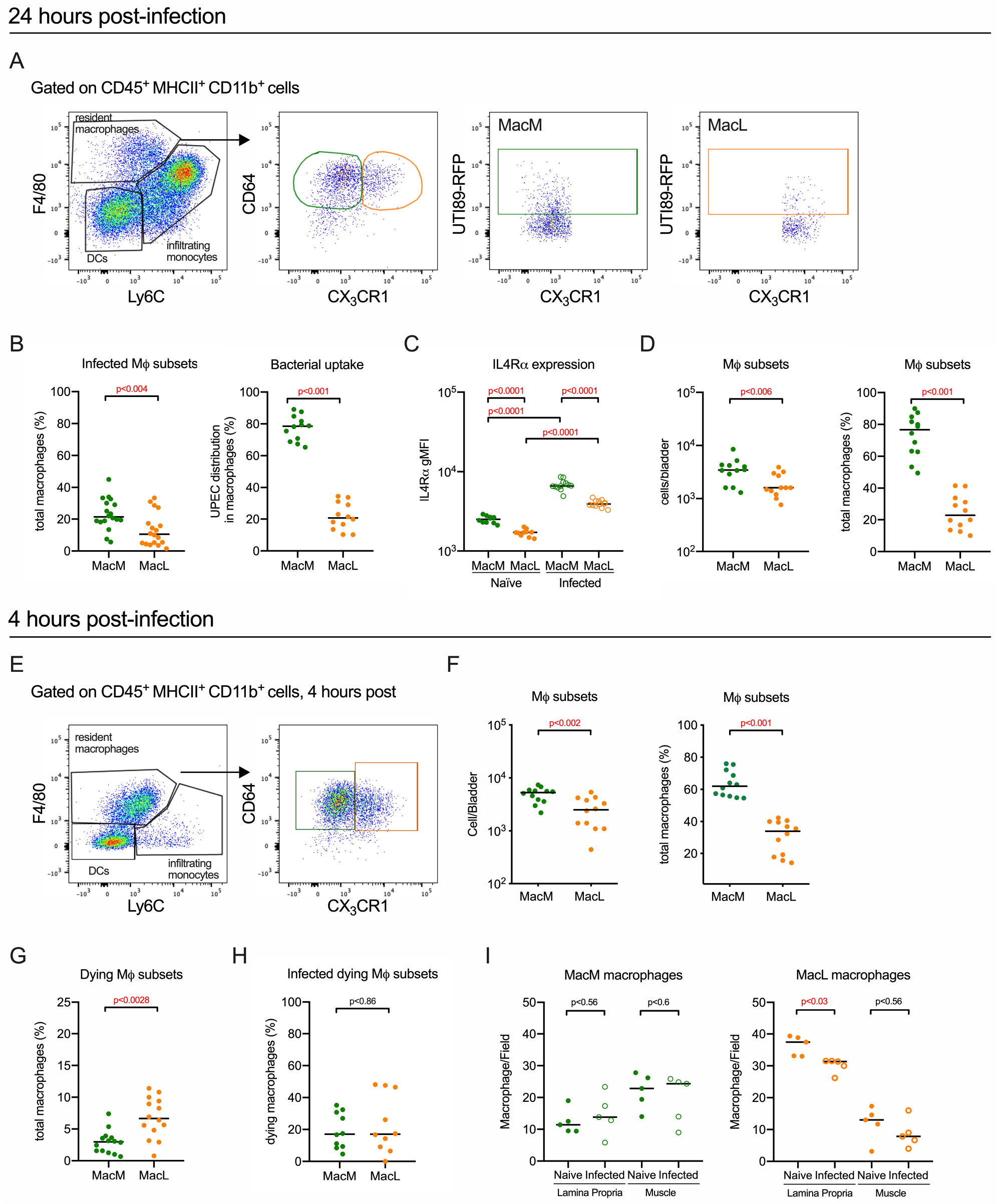
Macrophage subsets have divergent roles in response to UTI. See also Figure S1. (**A**-**H**)Female C57BL6/J were infected with 10^7^ CFU of UTI89-RFP and bladders were analyzed by flow-cytometry at (**A**-**D**) 24 hour or (**E**-**H**) 4 hours PI. (**A**) Dot plots show gating strategy to identify resident macrophage subsets and cells containing bacteria at 24 hours PI. (**B**) Graphs depict the percentage of macrophage subsets infected and the distribution of UPEC between the MacM and MacL subsets. (**C**) The graph shows the gMFI of IL-4Rα expression on macrophage subsets in naïve mice and at 24 hours PI. (**D**) Graphs show the total number and frequency of each macrophage subset in the bladder at 24 hours PI. (**E**) Dot plots show gating strategy to identify macrophage subsets at 4 hours PI. (**F**) Graphs show the total number and frequency of each macrophage subset in the bladder at 4 hours PI. (**G-H**) Graphs depict the percentage of macrophage subsets that were (**G**) labeled with a live/dead marker and the proportion of those dying macrophages that contained UPEC at 4 hours PI. (**I**) Graphs show the quantification of MacM and MacL macrophages by immunofluorescence, in the lamina propria and muscle of naïve mice (filled circles) and at 4 hours PI (open circles). In **B-D**, **F**-**H**, data are pooled from 3 experiments, n=3-6 mice per experiment. In **I**, data are pooled from 2 experiments, n=2-3 mice per experiment. Each dot represents one mouse, lines are medians. In **D**, **F**, significance was determined using the nonparametric Mann-Whitney test to compare the numbers of macrophages subsets and the nonparametric Wilcoxon matched-pairs signed rank test to compare the percentages of each macrophage subset. In **B-C**, and **G**-**I** significance was determined using the nonparametric Mann-Whitney test and resulting p-values were corrected for multiple testing using the false discovery rate (FDR) method. All calculated/corrected p-values are shown and p-values meeting the criteria for statistical significance (p <0.05) are depicted in red.

In the course of our studies, we observed that the total number and proportion of MacL macrophages was significantly lower than MacM macrophages at 24 hours PI, whereas in naïve mice, both the number and proportion of the macrophage subsets were equivalent (**Figure 4D, Figure 1B**). To rule out the contribution of differentiated monocyte-derived cells to the macrophage pool, we assessed total macrophage cell numbers in the bladder at 4h PI, when there is minimal infiltration of monocytes (**Figure 4E**) (Mora-Bau et al., 2015). Interestingly, macrophage subset numbers and proportions were significantly different at 4 hours PI, as well (**Figure 4F**). As the total numbers of each subset were not increased over naïve levels (see **Figure 1B**), we hypothesized that macrophages die during infection, particularly as apoptosis pathways were more highly expressed in MacL macrophages. Indeed, at 4 hours PI, we found that a significantly higher proportion of MacL macrophages were dying compared to MacM macrophages using a cell viability dye, which labels dying/dead cells (**Figure 4G, Figure S1B**).As UPEC strains can induce macrophage death *in vitro* (Murthy et al., 2018; Schaale et al., 2016), we asked whether macrophage cell death was induced by UPEC *in vivo*. We observed that only 20% of dying or dead cells in each subset were infected (**Figure 4H**), suggesting that macrophage death was not primarily driven by UPEC uptake. To determine whether macrophage cell death was confined to a distinct location, we quantified macrophage subset numbers in the muscle and lamina propria. We observed that at 4h PI, only MacL macrophages located in the lamina propria were reduced in numbers compared to naïve mice (**Figure 4I**).Given that in the first hours after infection, the urothelium exfoliates massively (Mulvey et al., 1998), these results suggest that macrophage death, specifically in the lamina propria, may be due to the loss of a survival factor in this niche. Altogether, we functionally validated the divergent gene expression observed between macrophage subsets, in which MacM macrophages are more phagocytic and MacL macrophages are more prone to die, suggesting that gene expression differences translate to differing roles for these subsets in the response to infection.

### Monocyte-derived macrophages replace resident macrophage subsets after infection

As we observed macrophages dying during infection, we investigated the change in macrophage numbers over time as animals resolved their infection. Both populations significantly decreased at 24 hours PI, then subsequently increased in numbers at 7 days PI, and returned to numbers just above homeostatic levels at 4 weeks PI (**Figure 5A**). With the dynamic increase of macrophage numbers over the course of UTI, we hypothesized that infiltrating monocytes replace resident macrophage subsets during infection. To test this hypothesis, we used the CCR2^CreERT2^ Rosa26^tdTomato^ mouse, in which administration of 4OHT induces recombination in CCR2-expressing cells, such as circulating Ly6C^+^ monocytes, leading to irreversible labeling of these cells *in vivo* (Croxford et al., 2015). Importantly, blood monocytes and bladder resident macrophages are not tomato+ in untreated mice (**Figure S2**). We administered 4OHT to naïve mice, and then, 24 hours later, infected half of the treated mice with 10^7^ CFU of UTI89. At the same time as infection, we analyzed the labeling efficiency in circulating classical Ly6C^+^ monocytes, finding that approximately 80% of Ly6C^+^ monocytes were labeled in both naïve and infected mice (**Figure 5B**). After 6 weeks, when animals had resolved their infection, there were no labeled circulating Ly6C^+^ monocytes in naïve or post-infected mice (**Figure 5B**). When we analyzed the bladders of naïve mice six weeks after the 4OHT pulse, only 2.8% ± 0.8% of MacM and 2.2% ± 0.6% of MacL macrophage subsets were labeled, supporting that monocytes make minimal contributions to bladder macrophage subsets in the steady state (**Figure 5C**). At 6 weeks post-infection, the total numbers of macrophage subsets finally returned to homeostatic levels (**Figure 5D**), but post-infection MacM and MacL macrophages had more tomato+ cells (MacM 6.5% ± 3%, MacL 8.1% ± 8.9%) than their naïve counterparts. The number of labeled macrophages in tissue was dependent upon the efficiency of initial labeling in the circulation, and so to mitigate this bias, we normalized labeling in bladder macrophages at 6 weeks using the percentage of labeled circulating monocytes at the time of infection, finding that the normalized proportion of labeled macrophages was significantly increased in post-infected mice compared to naïve mice 6 weeks after 4OHT pulse (**Figure 5C**). These data support that monocytes infiltrate the bladder during infection, and may contribute to the return of macrophage subsets to homeostatic levels.

**Figure 5.**
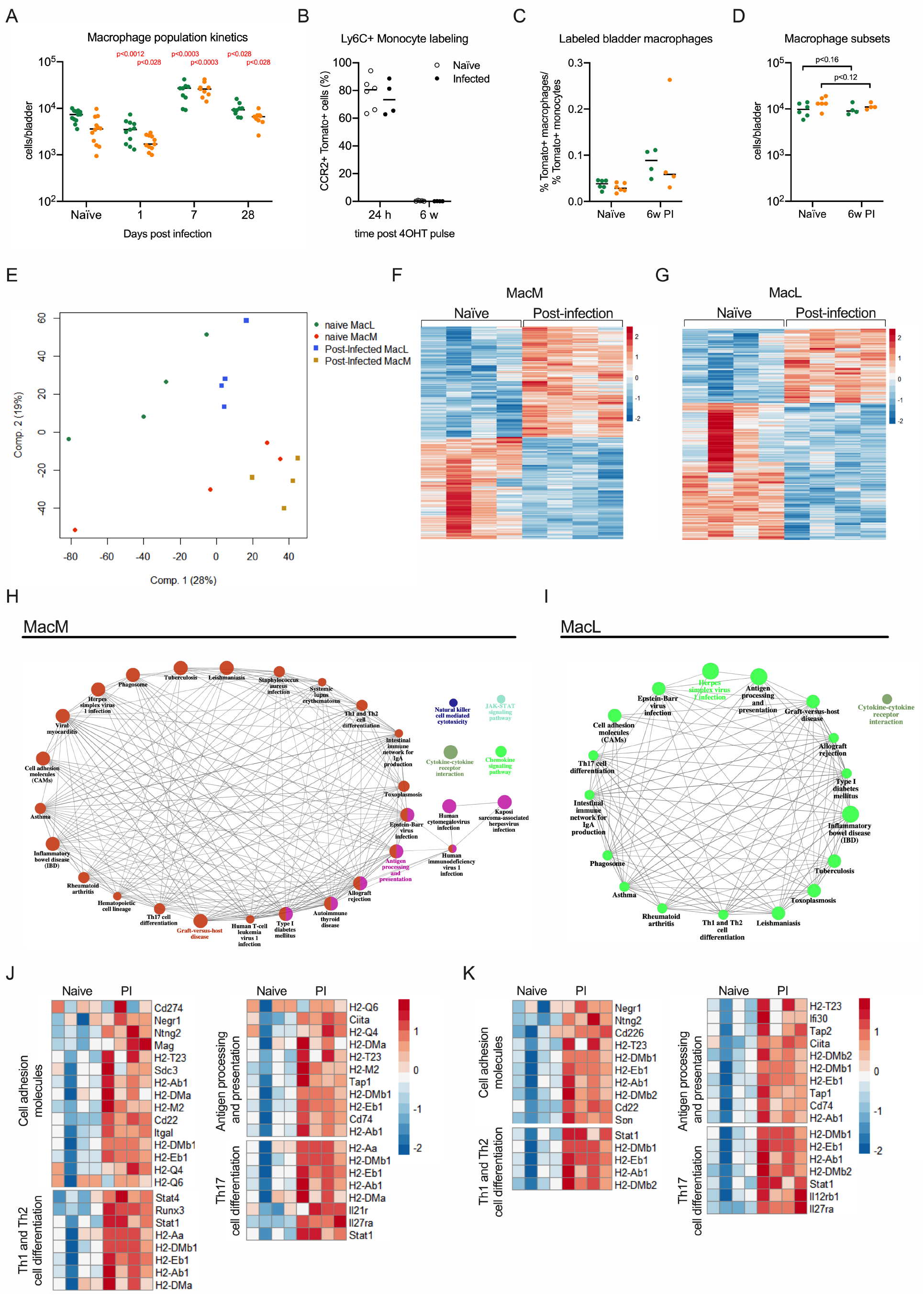
Macrophage subsets are supplemented by monocyte-derived cells after UTI and take on distinct transcriptional profiles. See also Figure S2. (**A**) The graph shows total number of MacM (green dots) and MacL (orange dots) in naïve mice and 1, 7, and 28 days PI. (**B-C**) Adult female CCR2^CreERT2^ Rosa26^tdTomato^ mice were pulsed with 4OHT and 24 hours later, half were infected with 10^7^ CFU of UTI89-RFP. (**B**) The graph shows the percentage of tomato+ Ly6C^+^ circulating monocytes at 24 hours and 6 weeks after the 4OHT pulse. (**C**) The graph depicts the percentage of tomato+ bladder macrophage subsets at 6 weeks post 4OHT pulse, normalized by the percentage of tomato+ Ly6C^+^ monocytes at 24 hours post 4OHT pulse. (**D**) The graph shows the total number of macrophage subsets in naïve mice and at 6 weeks post-infection. (**E**) Principle component analysis of all DEG from naïve and post-infected bladder macrophage subsets. (**F**-**G**) The heat maps show the differentially expressed genes between naïve and 6 week post-infected (**F**) MacM (513 genes) and (**G**) MacL (617 genes) subsets. (**H**-**I**) Using KEGG pathway analysis of significantly up-regulated genes, depicted are pathways enriched in 6 week post-infected (**H**) MacM and (**I**) MacL macrophages. (**J**-**K**) Shown are up-regulated genes associated with selected pathways in 6 week post-infected (**J**) MacM and (**K**) MacL macrophages. In (**H**-**I**), The size of the nodes reflects the statistical significance of the term. (*Q*<0.05; terms >3 genes; % genes/term >3; *κ* 0.4). In **A, C,** and **D,** significance was determined using the nonparametric Mann-Whitney test, comparing the number of each subset during infection to the control naïve subset number and resulting p-values were corrected for multiple testing using the false discovery rate (FDR) method. For each day, the higher left-shifted p-value refers to MacM macrophages and the lower right shifted p-value refers to MacL macrophages. All calculated/corrected p-values are shown and p-values meeting the criteria for statistical significance (p <0.05) are depicted in red.

As monocytes generally have different origins and developmental programs compared to tissue-resident macrophages, we used RNA sequencing to determine whether the macrophage pool in post-infected bladders was different from naïve tissue resident cells. Using principle component analysis, we compared bladder macrophage subsets from 6 weeks post-infected mice to their naïve counterparts. We found that macrophages clustered more closely together by subset, rather than by infection status, or, in other words, naïve and post-infected MacL macrophages clustered more closely to each other than either sample clustered to naïve or post-infected MacM macrophages (**Figure 5E**). 513 genes (247 genes down-regulated and 266 genes up-regulated) were different between naïve and post-infected MacM macrophages (**Figure 5F**). 617 genes (401 genes down-regulated and 216 genes up-regulated) were differentially expressed between the naïve and post-infected MacL subset (**Figure 5G**). Applying gene set enrichment analysis to up-regulated genes in the post-infected macrophage subsets, we detected more common pathways between the subsets including enrichment of genes linked to pathways such as *antigen presentation*, *cell adhesion molecules*, *Th1, Th2 and Th17 cell differentiation*, and *chemokine signaling pathway* (**Figure 5H, I**). Although the enriched genes were not identical within each subset for these pathways, some common up-regulated genes included those encoding for histocompatibility class 2 molecules, such as *H2-Ab1, H2-Eb1, H2-DMb1, Ciita,* and the Stat1 transcription factor (**Figure 5I-J**). As differentiation of monocytes into macrophages includes up-regulation of cell adhesion and antigen presentation molecules (Gross-Vered et al., 2020), including in the bladder (Mora-Bau et al., 2015), these data further support that monocytes specifically contribute to the post-infection bladder-resident macrophage pool.

These results show that in the context of UTI, dying macrophages are replaced by monocyte-derived cells. Tissue resident macrophage subsets maintain their separate identities distinct from each other after infection, although each subset also takes on a different transcriptional profile compared to their naïve counterparts, with upregulated expression of genes related to adaptive immune responses.

### Macrophage depletion before challenge improves bacterial clearance

Given that post-infected macrophage subsets upregulated pathways linked to inflammatory diseases and the adaptive immune response, we hypothesized that one or both macrophage subsets would induce improved bacterial clearance to a challenge infection. To test this hypothesis, we infected mice with 10^7^ CFU of kanamycin-resistant UTI89-RFP. Four weeks later, when the infection was resolved, mice were challenged with 10^7^ CFU of the isogenic ampicillin-resistant UPEC strain, UTI89-GFP and bacterial burden measured at 24 hours PI. To test the contribution of the macrophage subsets to the response to challenge infection, we used different concentrations of anti-CSF1R depleting antibody to differently target the two macrophage subsets directly before challenge infection (**Figure 6A**, experimental scheme). Using 500 μg of anti-CSF1R antibody, we depleted 50% of MacM and 80% of MacL macrophages, whereas depletion following treatment with 800 μg of anti-CSF1R antibody reduced MacM macrophages by 80% and the MacL subset by more than 90% (**Figure 6B, Figure S3A**). Importantly, 24 hours after anti-CSF1R antibody treatment, the number of circulating neutrophils, eosinophils, NK, T, and B cells was not different from mock-treated mice at either concentration (**Figure S3B**). Classical Ly6C^+^ monocytes were modestly reduced in mice treated with 800μg of anti-CSF1R antibody, but were unchanged in mice receiving 500 μg of depleting antibody. Antibody treatment did not change circulating non-classical monocyte numbers (**Figure S3B**).After challenge infection, the bacterial burden was not different in mice treated with 500 μg of anti-CSF1R compared to mock-treated mice (**Figure 6B**). By contrast, mice depleted with 800 μg of anti-CSF1R had reduced bacterial burdens, indicative of a stronger response post-challenge compared to non-depleted mice (**Figure 6C**).

**Figure 6.**
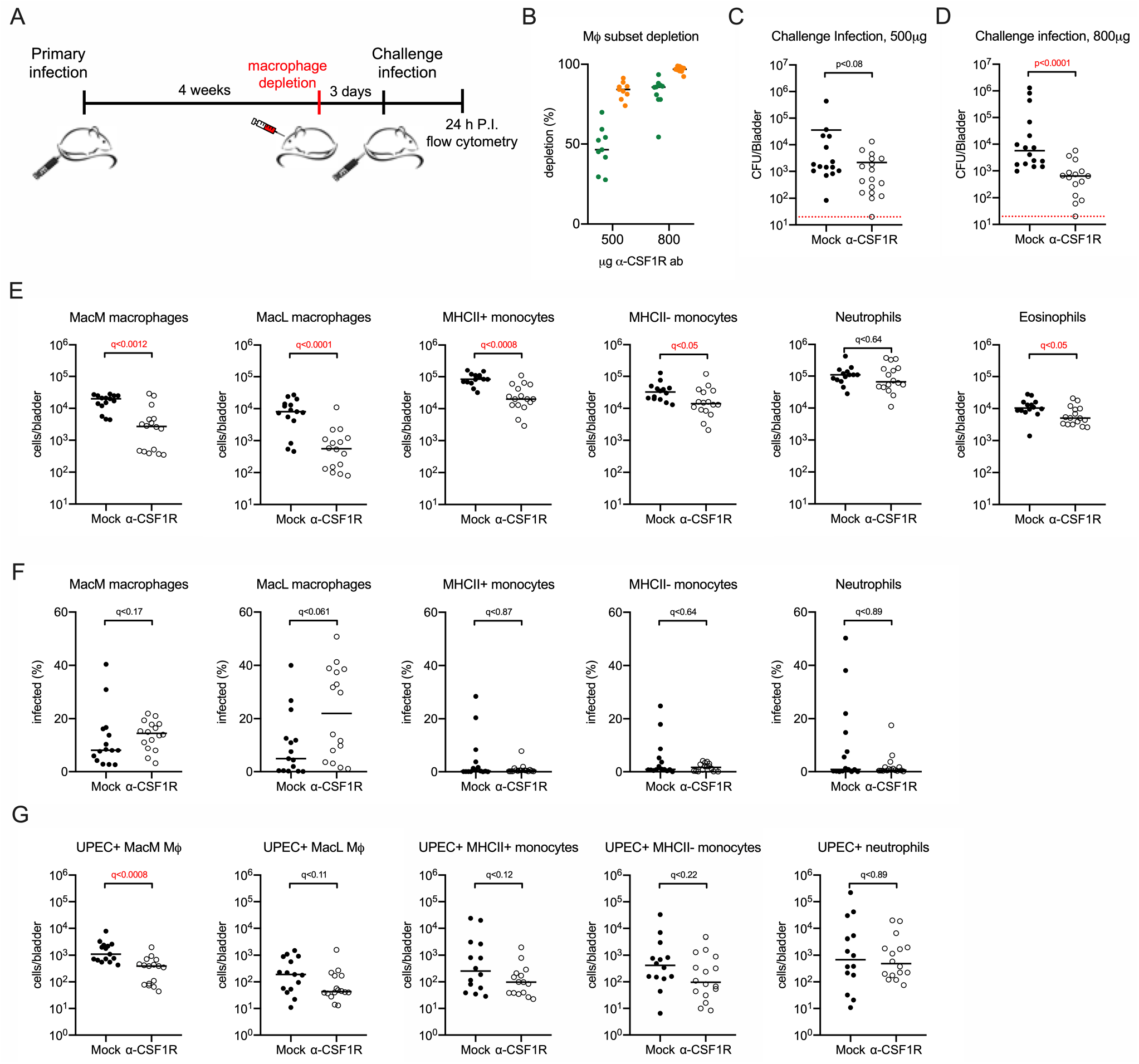
Macrophage depletion before challenge promotes improved bacterial clearance. See also Figure S3. (**B**) The graph shows the efficacy of macrophage subset depletion in naïve female C57BL/6 mice treated with 500 μg or 800 μg of anti-CSF1R antibody. (**C-D**) Graphs show bacterial burden per bladder 24 hours post-challenge in 7-8 week old female C57BL/6 mice that were infected with 10^7^ CFU of UTI89-RFP, allowed to resolve the infection over 4 weeks, then PBS (mock)-treated or treated with (**C**) 500 μg, or (**D**) 800 μg of anti-CSF1R antibody 24 hours prior to being challenged with 10^7^ CFU of the isogenic UPEC strain UTI89-GFP. (**E**-**G**) Mice were infected with 10^7^ CFU of UTI89-RFP, allowed to resolve the infection over 4 weeks, then PBS (mock)-treated or treated with 800 μg of anti-CSF1R antibody 24 hours prior to being challenged with 10^7^ CFU of the isogenic UPEC strain UTI89-GFP. Graphs depict the (**E**) total number of the indicated cell type, (**F**) the percentage of the indicated cell type that was infected, and (**G**) the total number of the indicated cell type that contained UPEC at 24 hours post-challenge in mice treated with PBS or 800 μg of anti-CSF1R antibody. Data are pooled from 3 experiments, n=3-6 mice per experiment. Each dot represents one mouse, line are medians. In **C-G**, significance was determined using the nonparametric Mann-Whitney test and resulting p-values were corrected for multiple testing using the false discovery rate (FDR) method. All calculated/corrected p-values are shown and p-values meeting the criteria for statistical significance (p <0.05) are depicted in red.

Neutrophils take up a majority of UPEC at early time points during UTI (Mora-Bau et al., 2015). Therefore, we hypothesized that the improved bacterial clearance in macrophage-depleted mice may be due to increased infiltration of inflammatory cells, such as neutrophils. At 24 hours post-challenge infection we observed that while the numbers of resident macrophage subsets, MHCII^+^ monocytes, and MHCII^−^ monocytes in macrophage-depleted mice were reduced compared to mock-treated mice, as expected, the numbers of infiltrating neutrophils were unchanged by antibody treatment (**Figure 6E**, **Figure S3C**, gating strategy). Interestingly, fewer eosinophils infiltrated the tissue in macrophage-depleted mice, although the impact of this is unclear as their role in infection is unknown (**Figure 6E**). Given that neutrophil infiltration was unchanged and that monocytes, which also take up a large number of bacteria during infection, were reduced in number, we considered that improved bacterial clearance in macrophage-depleted mice may be due to increased bacterial uptake on a per cell basis during challenge infection. However, bacterial uptake was not different between depleted and mock-treated mice in neutrophils, MHCII^+^ and MHCII^−^ monocytes, or either macrophage subset (**Figure 6F**).The lower numbers of the MacM subset in macrophage-depleted mice translated to lower numbers of infected MacM macrophages (**Figure 6E, F** respectively). However, we observed no differences in the numbers of infected MacL macrophages, neutrophils, MHCII^+^ or MHCII^−^ monocytes in macrophage-depleted mice compared to non-depleted animals (**Figure 6G**). Together, these results support that MacM macrophages negatively impact bacterial clearance in a challenge infection, but not at the level of direct bacterial uptake or myeloid cell infiltration.

### Depleting macrophages leads to a Th1 biased immune response during challenge infection

As infiltration of inflammatory cells or number of infected cells during challenge infection was not changed in macrophage-depleted mice, we questioned whether another host mechanism is involved in bacterial clearance. Exfoliation of infected urothelial cells is a host mechanism to eliminate bacteria (Anderson et al., 2003; Mulvey et al., 1998). We hypothesized that macrophage-depleted mice have increased urothelial exfoliation during challenge infection, leading to reduced bacterial numbers. We quantified the mean fluorescence intensity of uroplakins, proteins expressed by terminally differentiated urothelial cells (Wu et al., 1990), from bladders of post-challenged mice, depleted of macrophages or not (**Figure 7A**). We did not detect a significant difference in urothelial exfoliation between mock treated animals and mice depleted of macrophage prior to challenge infection, supporting that urothelial exfoliation is not the underlying mechanism behind improved bacterial clearance in macrophage-depleted mice (**Figure 7B**). Infiltration of inflammatory cells is associated with bladder tissue damage and increased bacterial burden (Ingersoll et al., 2008). As we observed fewer monocytes and eosinophils in macrophage-depleted mice during challenge infection, we investigated whether reduced cell infiltration was associated with less tissue damage. We assessed edema formation by quantifying area of the lamina propria in post-challenged bladders, depleted of macrophages or not (**Figure 7A**). Surprisingly, we did not detect a difference in edema formation between non-depleted mice and mice depleted of macrophage before challenge infection (**Figure 7C**).

**Figure 7.**
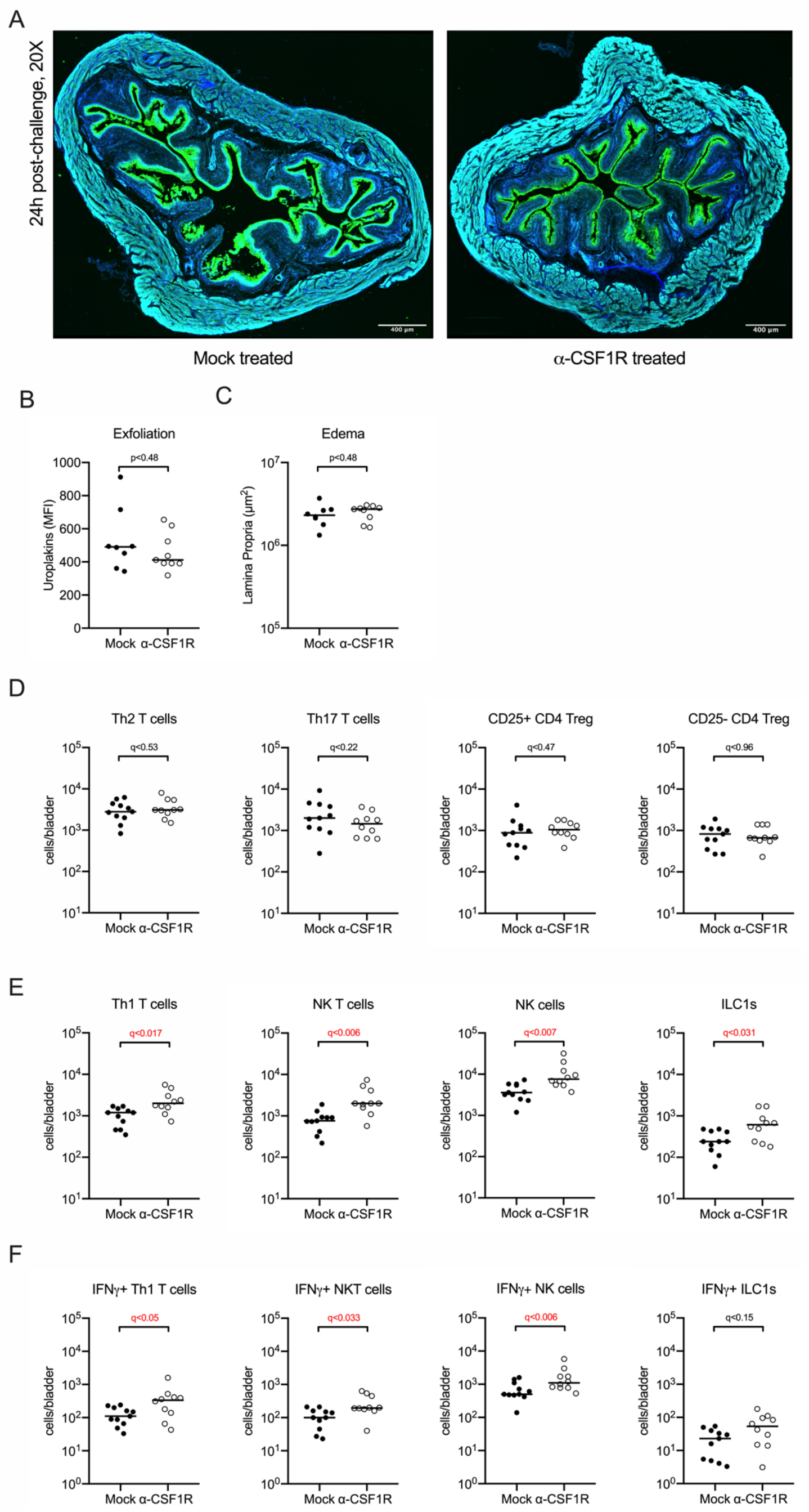
Depletion of replaced macrophages leads to a type I immune bias. See also Figure S4. 7-8 week old female C57BL/6 mice were infected with 10^7^ CFU of UTI89, allowed to resolve the infection over 4 weeks, then PBS (mock)-treated or treated with 800 μg of anti-CSF1R antibody 24 hours prior to being challenged with 10^7^ CFU of UTI89. (**A**) Representative confocal images of bladders from mice treated with PBS or 800 μg of anti-CSF1R antibody 24 hours post-challenge. Uroplakin – green, phalloidin – turquoise, DAPI – blue. (**B**) The graph shows the mean fluorescent intensity of uroplakin expression, quantified from imaging, at 24 hours post-challenge. (**C**) The graph shows the area of the lamina propria, quantified from imaging, at 24 hours post-challenge. (**D-F**) Graphs depict the (**D**-**E**) total number of the indicated cell type or (**F**) the total number of the indicated cell type expressing IFN-γ at 24h PI. Data are pooled from 2 experiments, n=4-6 mice per experiment. Each dot represents one mouse, line are medians. In **B**-**C** and **D**-**F**, significance was determined using the nonparametric Mann-Whitney test and resulting p-values were corrected for multiple testing using the false discovery rate (FDR) method. All calculated/corrected p-values are shown and p-values meeting the criteria for statistical significance (p <0.05) are depicted in red.

As we observed fewer eosinophils in macrophage-depleted mice during challenge infection, and our previous work demonstrated that type 2 immune response-related cytokines are expressed early in UTI (Zychlinsky Scharff et al., 2019), we assessed the polarity of the T cell response to challenge infection (gating strategy, **Figure S4**). Macrophage depletion did not alter the infiltration of T regulatory cells, or T_h_2 or T_h_17 T helper subsets (**Figure 7D**). However, macrophage depletion did correlate with an increase in the numbers of T_h_1 T cells, NKT cells, NK cells, and type 1 innate lymphoid cells (ILC1s) (**Figure 7E**).Additionally, in macrophage-depleted mice, T_h_1 T cells, NKT cells, and NK cells had higher IFN-γ production compared to mock-treated mice (**Figure 7F**), suggesting that in the absence of post-infected macrophages, a more pro-inflammatory, bactericidal response to challenge infection arises in the bladder.

## Discussion

Despite numerous studies of macrophage ontogeny and function in many organs, the developmental origin and role of bladder macrophages is largely unknown. Here, we investigated this poorly understood compartment in homeostasis and a highly inflammatory infectious disease, UTI. A homogeneous macrophage population of yolk-sac and HSC fetal origin seeds the developing bladder; however, the yolk-sac macrophage pool is ultimately replaced at some point after birth. After birth, two subsets, MacM and MacL, arise in the tissue, localizing to the muscle layer and the lamina propria, respectively. These subsets share similar developmental origin, in that they are primarily HSC-derived in adulthood with a slow turn over by Ly6C^+^ monocytes in the steady state. Their distinct transcriptomics demonstrate that they play different roles in the bladder, at least in the context of infection. The MacM subset is poised to take up bacteria or potentially, infected dying host cells, while polarizing towards a more alternatively activated profile during UTI. MacL macrophages express a profile with greater potential for the induction of inflammation, and whether due to direct consequences of this inflammation, or potentially due to loss of the urothelium, undergo pronounced cell death during UTI.

In adult animals, tissue-resident macrophages are a mix of embryonic and monocyte-derived macrophages, with the exception of brain microglia (Ginhoux et al., 2010; Gomez Perdiguero et al., 2015). The contributions from embryonic macrophages and circulating monocytes to the adult bladder macrophage compartment is similar to that of the lung and kidney (Epelman et al., 2014a; Schulz et al., 2012). Interestingly, although two macrophage subsets reside in the adult bladder, only a single LYVE1^+^CX_3_CR1^+^ macrophage population was identified in embryonic and newborn bladders. As the bladder is fully formed in newborn mice (Jost, 1989), it is unlikely that macrophage subsets arise to meet the needs of a new structure, as is the case for peritubular macrophages in the testis (Mossadegh-Keller et al., 2017). Rather, although all structures are present, embryonic or prenatal bladder tissue demands are likely distinct from postnatal tissue remodeling in very young mice or macrophage functions in an adult bladder as in the first weeks after birth, urothelial cells undergo increased proliferation to establish the three layers of urothelium in adult bladders (Cohen et al., 1988). As these adult tissue niches become fully mature, they may provide different growth or survival factors, driving functional macrophage specialization in discrete locations in the tissue.

In the lung, spleen, bone marrow, and liver, a subpopulation of pro-resolving macrophages are present that phagocytize blood-borne cellular material to maintain tissue homeostasis (A-Gonzalez et al., 2017; N et al., 2017). These macrophages express *Mrc1* (encoding for CD206), *CD163*, and *Timd4* (encoding TIM4) (A-Gonzalez et al., 2017; N et al., 2017). MacM macrophages likely represent this subpopulation in the bladder, as they expressed higher levels of genes associated with a pro-resolving phenotype, including the efferocytic receptor TIM4, CD206, and CD163. It is also possible that, similar to muscularis macrophages in the gut, MacM macrophages interact with neurons to control muscle contraction in the bladder and limit neuronal damage during infection (Matheis et al., 2020; Muller et al., 2014). By contrast, up-regulated pathways in the MacL subset, in combination with their localization under the urothelium suggest that, similar to intestinal macrophages, they may regulate T cell responses to bladder microbiota or support urothelial cell integrity (Morhardt et al., 2019; Scott et al., 2018).

Supporting a functional consequence of expression of genes associated with *complement*, *endocytosis*, and *phagosome* pathways in the MacM subset, this subset took up more bacteria during infection. This was somewhat surprising as this macrophage subset is located farther from the lumen and urothelium, where infection takes place. It is possible, although challenging to empirically demonstrate, that the MacM subset recognizes dying neutrophils, or even dying MacL macrophages, that have phagocytosed bacteria. Indeed, significant numbers of MacL macrophages died in the first hours following infection, which reflects their enriched apoptosis pathway. It is less likely that the bacteria induced macrophage death as only a small, and equivalent, proportion of both subsets were infected. Instead, MacL macrophage death may be an important step to initiate immune responses to UPEC. In the liver, Kupffer cell death by necroptosis during *Listeria monocytogenes* infection induces recruitment of monocytes, which in turn phagocytose bacteria (Bleriot et al., 2015). Here, macrophage depletion before challenge infection resulted in decreased infiltration of monocytes, likely due to diminished numbers of these cells in circulation, and fewer eosinophils, however, bacterial burden was also decreased. This suggests that macrophage-mediated immune cell recruitment is not their primary function in the bladder. Infiltration of inflammatory cells is not the only way macrophage cell death regulates infection, however. For instance, pyroptotic macrophages can entrap live bacteria and facilitate their elimination by neutrophils *in vivo* (Jorgensen et al., 2016). As MacM macrophages express genes regulating iron metabolism, limiting iron to UPEC would also be a plausible mechanism to control bacterial growth (Robinson et al., 2018).

In the steady state, tissue-resident macrophages can self-maintain locally by proliferation, with minimal input of circulating monocytes (Hashimoto et al., 2013; Merad et al., 2002). By contrast, under inflammatory conditions, resident macrophages are often replaced by monocyte-derived macrophages (Aegerter et al., 2020; Bleriot et al., 2015; Merad et al., 2002; Misharin et al., 2017). Monocytes will differentiate into self-renewing functional macrophages if the endogenous tissue resident macrophages are depleted or are absent (Scott et al., 2016; van de Laar et al., 2016). Our results show that UPEC infection induces sufficient inflammation to foster newly recruited monocyte differentiation. It is likely, however, that more macrophage replacement occurs than we actually measured, as 4OHT treatment in CCR2^CreERT2^ Rosa26^tdTomato^ mice only labeled monocytes in the first 24 hours of infection.

Recruited monocyte-derived macrophages can behave differently than resident macrophages when activated, such as in the lung. Gamma herpes virus induces alveolar macrophage replacement by “regulatory” monocytes expressing higher levels of Sca-1 and MHC II (Machiels et al., 2017). These post-infected mice have reduced perivascular and peribronchial inflammation and inflammatory cytokines, and fewer eosinophils compared to mock-infected mice when exposed to house dust mite to induce allergic asthma (Machiels et al., 2017). Alveolar macrophages of mice infected with influenza virus are replaced by pro-inflammatory monocyte-derived macrophages. At 30 days post-infection, influenza-infected mice have more alveolar macrophages and increased production of IL-6 when challenged with *S. pneumoniae* compared to mock-infected mice, leading to less death (Aegerter et al., 2020). Although mechanisms regulating the phenotype of monocyte-derived macrophages are not known, the time of residency in the tissue and the nature of subsequent insults likely influence these cells. Indeed, the longer that recruited macrophages reside in tissue, the more similar they become to tissue-resident macrophages and no longer provide enhanced protection to subsequent tissue injury (Aegerter et al., 2020; Misharin et al., 2017). In contrast to these studies in the lung, we found that elimination of macrophages, including those recruited during primary infection, led to improved bacterial clearance during secondary challenge, although it is not clear what the long-term consequences on bladder homeostasis might be when a more inflammatory type 1 immune response arises during infection.

Overall, our results demonstrate that two unique subsets of macrophages reside in the bladder. During UTI, these cells respond differently, and a proportion of the population dies. Thus, a first UPEC infection induces replacement of resident macrophage subsets by monocyte-derived cells. When sufficient numbers of MacM macrophages, composed of resident and replaced cells, are depleted, improved bacterial clearance follows, suggesting a major role of this subset in directing the immune response to challenge infection. While these findings greatly improve our understanding of this important immune cell type, much remains to be uncovered, such as the signals and niches that contribute to the establishment of two subsets of bladder resident macrophages and their roles in the establishment and maintenance of homeostasis.

## Acknowledgements

We are thankful for insightful discussions, technical support, and critical reading of the manuscript by Florent Vermeulen. We gratefully acknowledge the Center for Translational Science (CRT)/Cytometry and Biomarkers Unit of Technology and Service (CB UTechS) at Institut Pasteur for support in conducting this study. We heartily thank the breeding team (DTPS-C2RA-Central Animal Facility platform) of the Institut Pasteur for support with breeding and maintaining our animals. We gratefully acknowledge the UTechS Photonic BioImaging (Imagopole), C2RT, Institut Pasteur, supported by the French National Research Agency (France BioImaging; ANR-10–INSB–04; Investments for the Future) as well as the Image Analysis Hub of the Institut Pasteur for support in conducting this study.

LLM is part of the Pasteur-Paris University (PPU) International PhD Program, which received funding from the European Union’s Horizon 2020 research and innovation program under the Marie Sklodowska-Curie grant agreement no. 665807 and from the Labex Milieu Intérieur (ANR-10-LABX-69-01). MB is supported by funding from the *Agence Nationale de la Recherché* (French National Research Agency) ANR-19-CE15-0015. EGP is supported by funding from the Revive Labex (ANR-10-LABX-73), the Fondation Schlumberger (FRM FSER 2017) and the Emergence(s) program from Ville de Paris (2016 DAE 190). MAI is supported by funding from the *Agence Nationale de la Recherché* (French National Research Agency) ANR-17-CE17-0014 and ANR-19-CE15-0015.

## Author contributions

Conceptualization: LLM and MAI; Methodology: LLM, MR, HV, RL RG, JSC, MB, EGP, MAI; Investigation and data analysis: LLM, MR, HV, RL RG, JSC, MB, EGP, MAI; Writing-Original Draft: LLM and MAI; Writing-Review & Editing: LLM, MR, HV, RL, RG, JSC, MB, EGP, MAI; Funding Acquisition: MB and MAI; Supervision: MAI.

## Materials and Methods

### Resource Availability

#### Lead Contact

Further information and requests for resources and reagents should be directed to and will be fulfilled by the Lead Contact, Molly A. Ingersoll.

#### Materials Availability

This study did not generate new unique reagents.

#### Data and Code Availability

RNA sequencing data used for Figures 3 and 5 are deposited in the NCBI Gene Expression Omnibus under the accession number GEO: GSE147909

### Experimental Model and Subject Details

#### Mice

All animals used in this study had free access to standard laboratory chow and water at all times. We used female C57BL/6J mice between 7 and 8 weeks old from Charles River, France. Female CX_3_CR1^GFP/+^ mice between 7 and 8 weeks were bred in house. CX_3_CR1^GFP/GFP^ mice, used to maintain our hemizygous colony, were a kind gift from Fabrice Chretien (Institut Pasteur). *Cdh5*-CreERT2 Rosa26^tdTomato^ mice were crossed to CX_3_CR1^GFP^ mice, producing *Cdh5*-CreERT2.Rosa26^tdTomato^.CX_3_CR1^GFP^ mice at Centre d’Immunologie de Marseille-Luminy. In *Cdh5*-CreERT2.Rosa26^tdTomato^.CX_3_CR1^GFP^ mice, cells expressing the CX_3_CR1 receptor are constitutively GFP+, and treatment with 4OHT conditionally labels hemogenically active endothelial cells (Gentek et al., 2018). We used female and male *Cdh5*-CreERT2.Rosa26^tdTomato^.CX_3_CR1^GFP^ mice between 8 and 11 weeks, at E16.5, and newborns. Flt3^Cre^.Rosa26^YFP^ mice were a kind gift from Elisa Gomez-Perdiguero (Institut Pasteur). CCR2^−/−^ mice were a kind gift from Marc Lecuit (Institut Pasteur). CCR2creERT2^BB^ mice were a kind gift from Burkhard Becher (University of Zurich) via Sebastian Amigorena (Institut Curie). CCR2creERT2^BB^ male mice were crossed to Rosa26^tdTomato^ females to obtain CCR2creERT2^BB^-tdTomato mice at Institut Pasteur. We used female CCR2creERT2^BB^-tdTomato mice between 6 and 7 weeks. Mice were anesthetized by injection of 100 mg/kg ketamine and 5 mg/kg xylazine and sacrificed by carbon dioxide inhalation. Experiments were conducted at Institut Pasteur in accordance with approval of protocol number 2016–0010 and dha190501 by the *Comité d’éthique en expérimentation animale Paris Centre et Sud* (the ethics committee for animal experimentation), in application of the European Directive 2010/63 EU. Experiments with Cdh5-CreERT2 mice were performed in the laboratory of Marc Bajenoff, Centre d’Immunologie de Marseille-Luminy in accordance with national and regional guidelines under protocol number 5-01022012 following review and approval by the local animal ethics committee in Marseille, France.

### Method Details

#### Flow cytometry of bladder tissue and blood

Samples were acquired on a BD LSRFortessa using DIVA software (v8.0.1) and data were analyzed by FlowJo (Treestar) software (version 10.0). The analysis of bladder and blood was performed as described previously (Mora-Bau et al., 2015). Briefly, bladders were dissected and digested in buffer containing 0.34 U/mL of Liberase in PBS at 37°C for one hour with manual agitation every 15 minutes. Digestion was stopped by adding PBS supplemented with 2% FBS and 0.2 μM EDTA (FACS buffer). Fc receptors in single cell suspensions were blocked with anti-mouse CD16/CD32 and stained with antibodies. Total cell counts were determined by addition of AccuCheck counting beads to a known volume of sample after staining, just prior to cytometer acquisition. To determine cell populations in the circulation, whole blood was incubated with BD PharmLyse, and stained with antibodies. Total cell counts were determined by the addition of AccuCheck counting beads to 10 μL of whole blood in 1-step Fix/Lyse Solution.

For intracellular staining, single cell suspensions were resuspended in 1 mL of Golgi stop protein transport inhibitor, diluted (1:1500) in RPMI with 10% FBS, 1% sodium pyruvate, 1X HEPES, 1X nonessential amino acid, 1% penicillin/streptomycin, 50 ng/mL of phorbol 12-myristate 13-acetate (PMA), and 1 μg/mL of ionomycin, and incubated for 4h at 37°C. Samples were washed once with FACS buffer, and Fc receptors blocked with anti-mouse CD16/CD32. Samples were stained with antibodies against surface markers and fixed and permeabilized with 1X fixation and permeabilization buffer and incubated at 4°C for 40-50 minutes protected from light. After incubation, samples were washed two times with 1X permeabilization and wash buffer from the transcription factor buffer kit and stained with antibodies against IFN-γ and the transcriptional factors RORγT, GATA3, T-bet, and FoxP3, diluted in 1X permeabilization and wash buffer at 4°C for 40-50 minutes protected from light. Finally, samples were washed two times with 1X permeabilization and wash buffer and resuspended in FACS buffer. Total cell counts were determined by addition of counting beads to a known volume of sample after staining, just prior to cytometer acquisition.

#### Histological and immunostaining for confocal microscopy

Whole bladders were fixed with 4% PFA in PBS for 1 hour, and subsequently washed with PBS. Samples were then dehydrated in 30% sucrose in PBS for 24 hours. Samples were cut transversally and embedded in optimal cutting temperature compound (OCT), frozen, and sectioned at 30 μm. Sections were blocked for 1 hour with blocking buffer (3% bovine serum albumin (BSA) + 0.1% Triton + donkey serum (1:20) in PBS), and washed three times. Immunostaining was performed using F4/80, LYVE1 antibodies, or polyclonal “asymmetrical unit membrane” antibodies, recognizing uroplakins (kind gift from Xue-Ru Wu, NYU School of Medicine, (Wu et al., 1990)) (1:200) in staining buffer (0.5% BSA + 0.1% Triton in PBS) overnight. Sections were washed and stained with phalloidin (1:350) and secondary antibodies (1:2000) in staining buffer for 4 hours. Finally, sections were washed and stained with DAPI. Confocal images were acquired on a Leica SP8 confocal microscope. Final image processing was done using Fiji (version 2.0.0-rc-69/1.52p) and Icy software (v1.8.6.0).

#### Timed pregnancies and in utero tamoxifen administration

Fate mapping of Cdh5-CreERT2 mice was performed as described previously (Gentek et al., 2018). Briefly, for reporter recombination in offspring, a single dose of 4-Hydroxytamoxifen (4OHT) supplemented with progesterone (1.2mg 4OHT and 0.6mg progesterone) was delivered by intraperitoneal injection to pregnant females at E7.5 or E10.5. Progesterone was used to counteract adverse effects of 4OHT on pregnancies. To fate map cells in CCR2creERT2^BB^-tdTomato mice, a single dose (37.5 μg/g) of 4OHT injection was delivered intraperitoneally.

#### Bone marrow chimeras

For shielded irradiation, 7-8 weeks old wildtype female CD45.1 or CD45.2 C57BL6/J mice were anesthetized and dressed in a lab-made lead diaper, which selectively exposed their tail, legs, torso, and head to irradiation, but protected the lower abdomen, including the bladder. Mice were irradiated with 9 Gy from an Xstrahl X-Ray generator (250kV 12mA), and reconstituted with ~3-4 x10^7^ bone marrow cells isolated from congenic (CD45.1) wildtype mice or CD45.2 CCR2^−/−^ mice.

#### RNA sequencing and bioinformatic analyses

Samples were obtained from the whole bladders of naïve and 6 week post-infected female C57BL/6J mice. Four separate fluorescence-activated cell sorts (FACS) were performed to generate biological replicates, each sort was a pool of 10 mouse bladders. Macrophage subsets were FACS purified into 350 μl of RLT plus buffer from the RNAeasy micro kit plus (1:100) β-mercaptoethanol. Total RNA was extracted using the RNAeasy micro kit following the manufacturer’s instructions. Directional libraries were prepared using the Smarter Stranded Total RNA-Seq kit Pico Input Mammalian following manufacturer’s instructions. The quality of libraries was assessed with the DNA-1000 kit on a 2100 Bioanalyzer and quantification was performed with Quant-It assays on a Qubit 3.0 fluorometer. Clusters were generated for the resulting libraries with Illumina HiSeq SR Cluster Kit v4 reagents. Sequencing was performed with the Illumina HiSeq 2500 system and HiSeq SBS kit v4 reagents. Runs were carried out over 65 cycles, including seven indexing cycles, to obtain 65-bp single-end reads. Sequencing data were processed with Illumina Pipeline software (Casava version 1.9). Reads were cleaned of adapter sequences and low-quality sequences using cutadapt version 1.11. Only sequences of at least 25 nucleotides in length were considered for further analysis and the 5 first bases were trimmed following library manufacturer’s instructions. STAR version 2.5.0a (Dobin et al., 2013), with default parameters, was used for alignment on the reference genome (Mus musculus GRCm38_87 from Ensembl version 87). Genes were counted using featureCounts version 1.4.6-p3 (Liao et al., 2014) from Subreads package (parameters: -t exon -g gene_id -s 1). Count data were analyzed using R version 3.4.3 and the Bioconductor package DESeq2 version 1.18.1 (Love et al., 2014). The normalization and dispersion estimation were performed with DESeq2 using the default parameters and statistical tests for differential expression were performed applying the independent filtering algorithm. A generalized linear model was set to test for the differential expression among the four biological conditions. For each pairwise comparison, raw p-values were adjusted for multiple testing according to the Benjamini and Hochberg (BH) procedure and genes with an adjusted p-value lower than 0.05 were considered differentially expressed. Count data were transformed using Variance Stabilizing Transformation (VST) to perform samples clustering and PCA plot.

To perform pathway analysis, gene lists of differentially expressed genes were imported in the Cytoscape software (version 3.7.2), and analyses were performed using the ClueGO application with the Kyoto Encyclopedia of Genes and Genomes (KEGG) as the database. Significant pathways were selected using the threshold criteria *Q*<0.05; terms >3 genes; % Genes/term >3; *κ* 0.4.

#### Bacterial strains and infection

We used the human UPEC cystitis isolate UTI89 engineered to express the fluorescent proteins RFP or GFP and antibiotic resistant cassettes to either kanamycin (UPEC-RFP) or ampicillin (UPEC-GFP) to infect animals for flow cytometric and bacterial burden analyses (Mora-Bau et al., 2015). We used the non-fluorescent parental strain UTI89 for confocal imaging experiments and flow cytometric experiments with CCR2^CreERT2^ Rosa26^tdTomato^ mice (Mulvey et al., 2001). To allow expression of type 1 pili, necessary for infection (Hultgren et al., 1986), bacteria cultures were grown statically in Luria-Bertani broth medium for 18 hours at 37°C in the presence of antibiotics (kanamycin 50 μg/mL or ampicillin 100 μg/mL). Primary and challenge urinary tract infection were induced in mice as previously described (Mora-Bau et al., 2015; Zychlinsky Scharff et al., 2017). For challenge infection, urine was collected twice a week, for 4 weeks, to follow presence of bacteria in the urine. Once there were no UTI89-RFP bacteria in the urine, mice were challenged with UTI89-GFP bacteria, and sacrificed 24 hours post-challenge infection. To calculate CFU, bladders were aseptically removed and homogenized in 1 mL of PBS. Serial dilutions were plated on LB agar plates with antibiotics, as required.

#### Immune cell depletion

To produce anti-CSF1R antibody, the hybridoma cell line AFS98 (kind gift from Miriam Merad at Icahn School of Medicine at Mount Sinai) (Hashimoto et al., 2011) was cultured in a disposable reactor cell culture flasks for 14 days, and antibodies were purified with disposable PD10 desalting columns. To deplete macrophages, wildtype C57BL/6 mice received intravenous (I.V.) injection of anti-CSF1R antibody (2 mg/mL) diluted in PBS. Animals received two or three I.V. injections, on consecutive days, of anti-CSF1R antibody or PBS. To deplete macrophages with a final concentration of 500 μg of anti-CSF1R, we administered 250 μg/mouse on day 1, and 250 μg/mouse on day 2. To deplete macrophages with a final concentration of 800 μg of anti-CSF1R, we administered 400 μg/mouse on day 1, 200 μg/mouse on day 2, and 200 μg/mouse on day 3 to minimize the impact on circulating monocytes.

### Quantification and Statistical Analysis

#### Quantification of macrophage subsets, urothelial exfoliation, and tissue edema

To quantify the number of macrophage subsets in bladder tissue, 6 to 7 images were randomly acquired of each of the areas of the muscle and lamina propria per mouse in wildtype C57BL/6 female mice with 40X magnification in a SP8 Leica microscope. Maximum intensity Z-projections were performed, and individual macrophage subsets were counted using Icy software (v1.8.6.0). To quantify urothelial exfoliation and tissue edema, images from whole bladder cross-sections were acquired using 20X magnification in a SP8 Leica microscope. Maximum intensity Z-projections were performed, the urothelium was delimited and mean fluorescence intensity of uroplakin staining was measured using Fiji (v1.51j) software. To quantify tissue edema, the lamina propria was delimited and the area was measured using Fiji software (v1.51j).

#### Statistical analysis

Statistical analysis was performed in GraphPad Prism 8 (GraphPad, USA) for Mac OS X applying the non-parametric Wilcoxon test for paired data or the non-parametric Mann-Whitney test for unpaired data in the case of two group comparisons. In the case that more than 2 groups were being compared or to correct for comparisons made within an entire analysis or experiment, calculated *p*-values were corrected for multiple testing with the false discovery rate (FDR) method, (https://jboussier.shinyapps.io/MultipleTesting/), to determine the false discovery rate adjusted *p*-value. All calculated *p*-values are shown in the figures, and those that met the criteria for statistical significance (*p* <0.05) are denoted with red text.

## Supplemental Information

**Figure S1, related to Figure 4.**
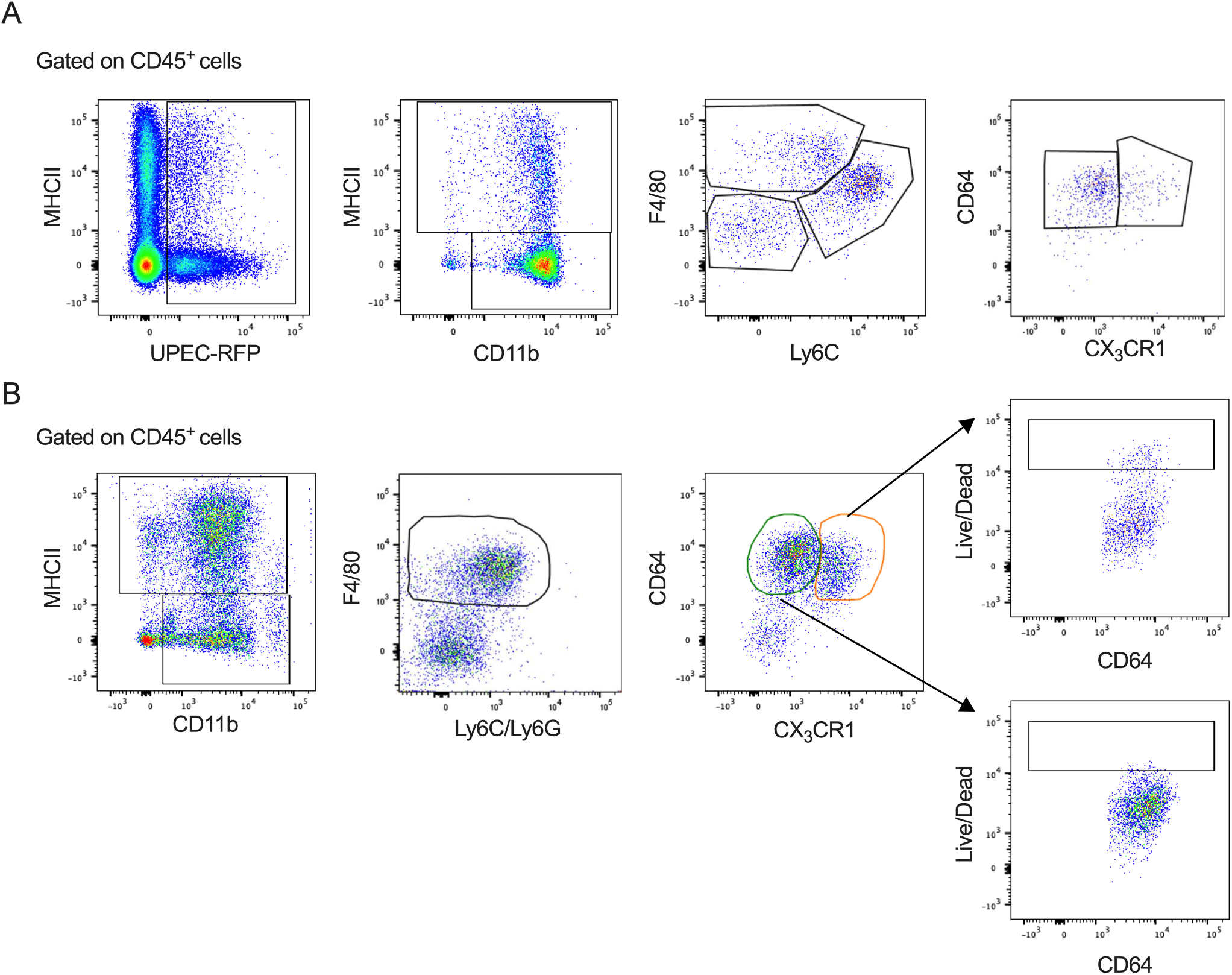
Gating strategy to identify macrophages containing bacteria. 7-8 week old female C57BL6/J mice were infected with 10^7^ CFU of UTI89-RFP and bladders were analyzed by flow-cytometry. (**A**) Dot plots show gating strategy to identify UPEC distribution in macrophage subsets at 24 hours PI. (**B**) Dot plots show gating strategy to identify macrophage subsets labeled with live/dead marker at 4 hours PI.

**Figure S2, related to Figure 5.**
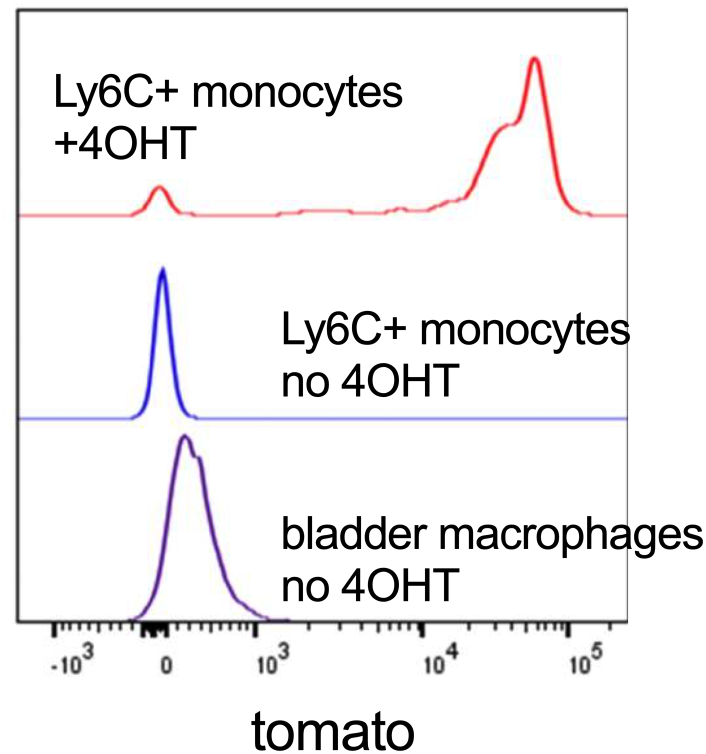
Blood monocytes and bladder resident macrophages are not tomato+ in untreated mice. Representative histogram shows tomato labeling in (top to bottom) Ly6C+ monocytes from a CCR2^CreERT2^ Rosa26^tdTomato^ mouse pulsed with 4OHT, Ly6C+ monocytes from a CCR2^CreERT2^ Rosa26^tdTomato^ mouse untreated, and total bladder resident macrophages from a CCR2^CreERT2^ Rosa26^tdTomato^ mouse.

**Figure S3, related to Figure 6.**
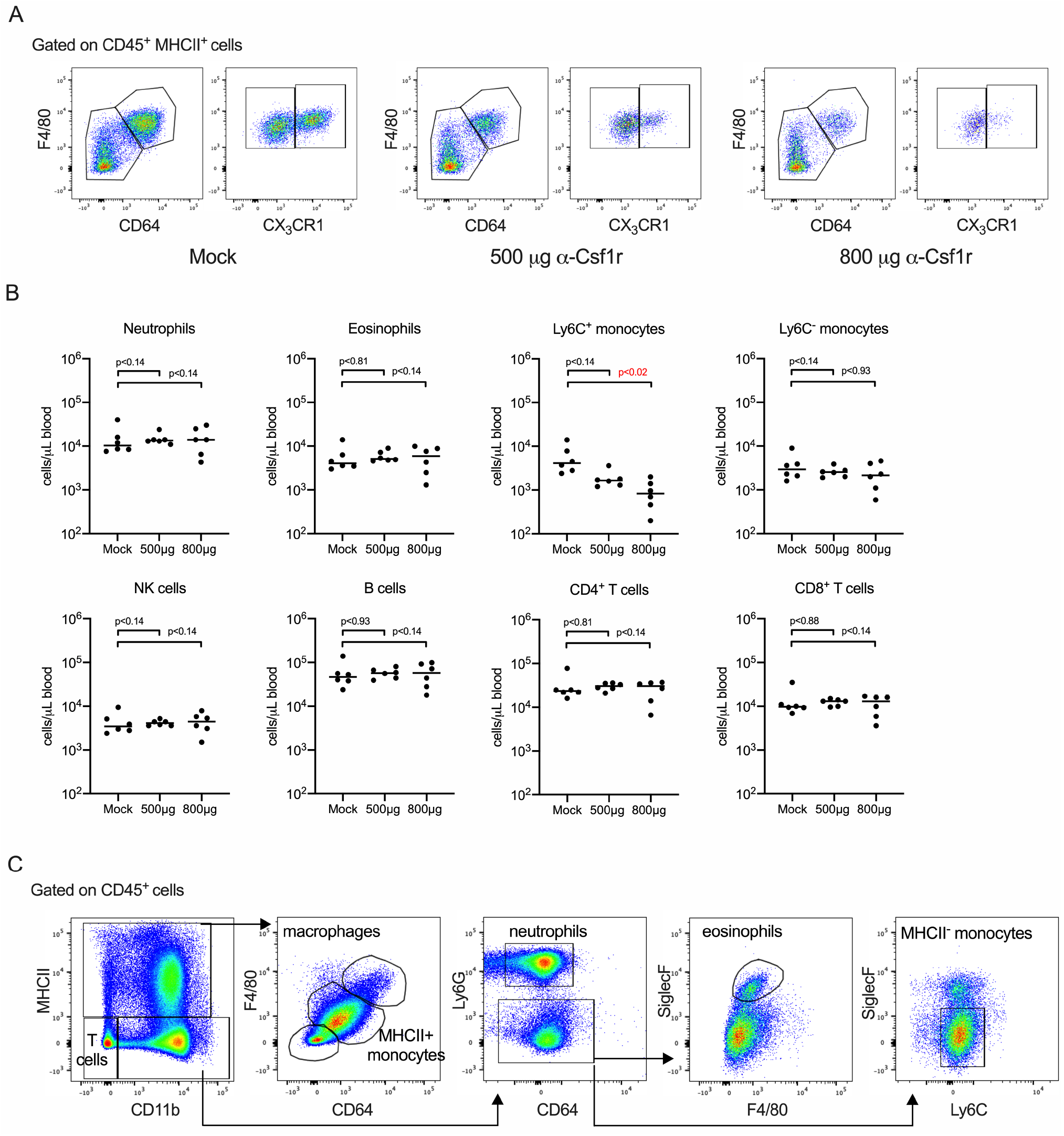
Macrophage depletion reduces circulating monocytes. Naïve female C57BL/6 mice were treated with PBS (mock), 500 μg, or 800 μg of anti-CSF1R antibody. (**A**) Representative dot plots depict efficacy of depletion of macrophage subsets in indicated scenarios. (**B**) Dot plots show the gating strategy used to identify macrophages, MHCII^+^ monocytes, MHCII^−^ monocytes, neutrophils, eosinophils, and T cells in the bladder at 24 hours PI. (**C**) Graphs depict the total number of the indicated cell types in circulation.

**Figure S4, related to Figure 7.**
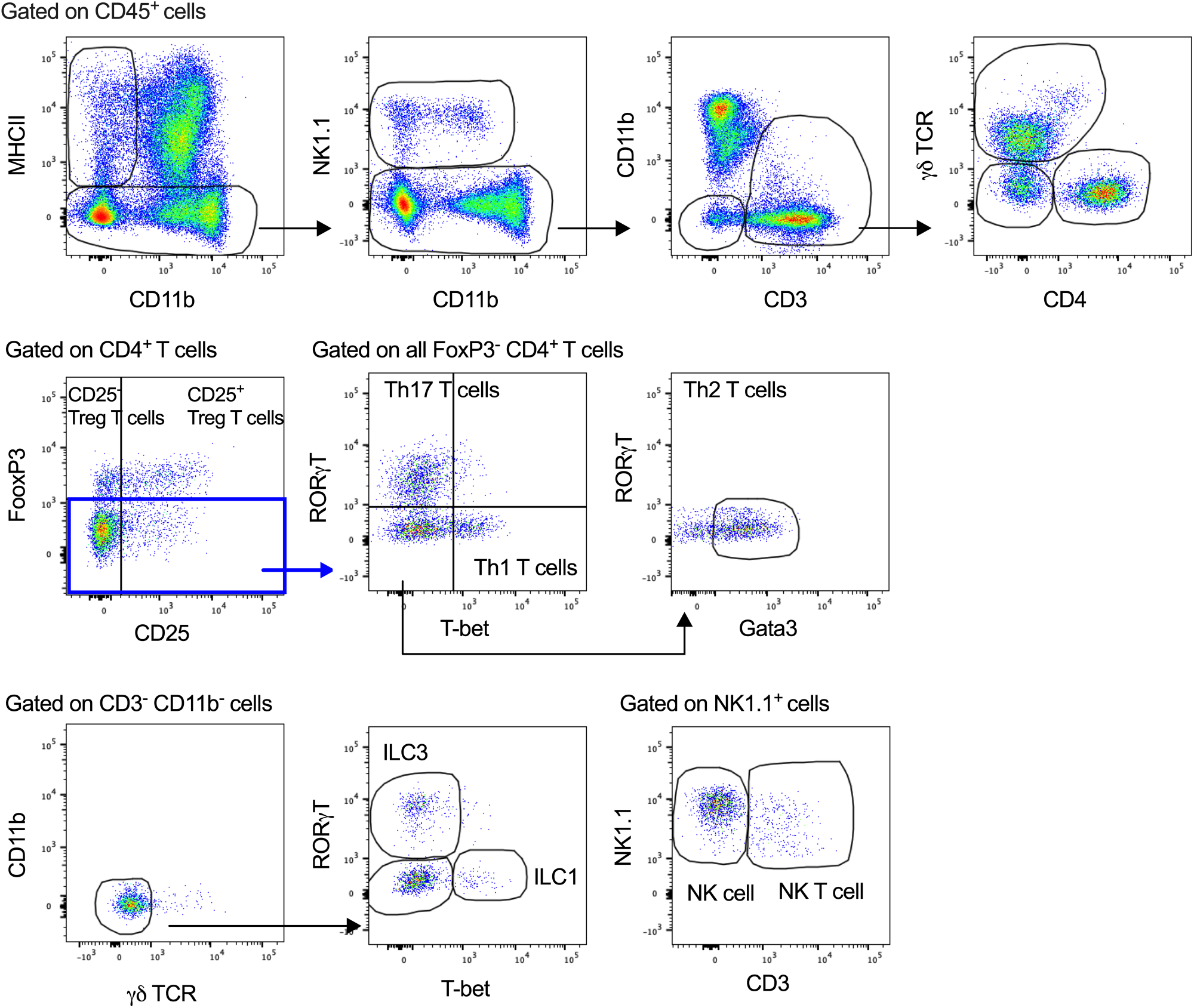
Gating strategy to identify T, NKT, NK and ILC cells. Dot plots show the gating strategy to identify T cell populations, NK T cells, NK cells, and innate lymphoid cells in the bladder at 24h post-challenge.

